# Hierarchical priors enable neural prediction of perceived biological motion

**DOI:** 10.1101/2025.10.09.681210

**Authors:** Ingmar E.J. de Vries, Floris P. de Lange, Moritz F. Wurm

## Abstract

Biological motion perception is an essential skill that allows us to quickly infer how others move. While this is often cast as inherently predictive process, it remains unclear to what extent different priors shape neural processing of biological motion. We investigated this by comparing the dynamic representational geometry of human magnetoencephalography (MEG) activity and observed biological motion stimuli by means of dynamic representational similarity analysis (dRSA). Under normal viewing conditions, neural representations were indeed predictive and followed an inverse hierarchy: high-level viewpoint-invariant body motion representations were visible before representations of viewpoint-dependent body motion and low-level visual features. Disrupting holistic priors by turning videos upside down selectively reduced high-level predictions. Instead, disrupting kinematics priors by temporal piecewise scrambling eliminated all motion prediction, with neural activity merely reacting to, rather than predicting, visual input. These findings reveal how low- and high-level priors jointly shape predictive neural processing of observed biological motion.

**Impact statement:** Disrupting holistic priors selectively interferes with neural prediction of observed biological motion

## Introduction

Biological motion perception is an essential skill for adaptive behavior such as social interaction (*1*, *2*), as it provides important social cues. This ability relies on our internal model of other agents and how they move (*3–6*). Our internal model generates probabilistic distributions, or priors, of the most probable external states and state changes, which in turn helps us infer the most probable causes of noisy and ambiguous sensory input. They include holistic priors of configural relations among joints or body parts, global body structure, body form templates, and biomechanical and gravity constraints (e.g., an upright body is more probable than an up-down inverted one, certain joint correlations occur more often (*7–14*)), and kinematics priors about biological motion continuity (e.g., smooth motion continuation is more probable, slower motions are more probable (*15–17*)). Importantly, for behavior to be adaptive and timely, our brain needs to continuously predict sensory input such as biological motion (*18–26*). While there is evidence that the prediction of simple sensory input relies on a combination of our priors and current sensory input (*27–29*), it remains unclear whether biological motion prediction relies on our holistic and kinematics priors of other agents and how they move, or whether general purpose motion prediction mechanisms such as extrapolation are sufficient. Based on the former hypothesis, we would expect our priors to activate predictive neural representations of observed biological motion. However, most empirical evidence for neural prediction comes from artificial paradigms that use simple isolated (often static) stimuli (*30–32*), and that only capture a single snapshot or the consequence of prediction rather than the continuous predictions themselves (*33*, *34*). They evidence that the brain *can* predict but preclude testing to what extent complex naturalistic input such as biological motion is predictively represented, whether such neural prediction relies on our priors, and how violations of these priors alter our brain’s representations.

We recently developed a novel dynamic extension to representational similarity analysis (dRSA) that quantifies exactly *what features* our brain represents *when* while viewing continuous and rich naturalistic input such as movies (*35*). Traditional RSA captures the similarity between neural representations and static stimulus models to infer whether the stimulus feature captured by the model is represented in the brain (*36*, *37*). Dynamic RSA extends this to temporally variable models to capture the representational dynamics of unfolding events along a feature dimension of interest. In short, it quantifies the strength of the match between continuous representations in the brain and continuous features of naturalistic input, across several hierarchical processing levels concurrently. Besides strength of representation, it also quantifies at millisecond-precision latency how neural representations temporally relate to (follow or precede) events of interest. Using this framework, we previously found an inverted temporal hierarchy of predictive neural representations of future movements during biological motion perception, such that high-level movement features were predicted *before* lower-level movement features (*35*). While these results demonstrate biological motion to be represented mostly predictively, rather than bottom-up (*35*), it remains unclear to what extent these hierarchical predictive representations rely on high-level priors about bodies and how they move, or rather on general motion extrapolation mechanisms.

Here we apply dRSA in the context of complex continuous human biological motion sequences (i.e., ballet dancing videos) to test if and how neural prediction relies on our priors. Specifically, we disrupted holistic priors of body form and structure, and action structures, as well as gravity priors, by showing the videos up-down inverted. This creates a perceptual experience that is very different from our typical experience of biological motion, and is thereby an effective way to impede biological motion perception and its associated neural processing, i.e., the so-called inversion effect (*6*, *11*, *38–41*). Indeed, akin to the inversion effect in face (*42*, *43*) and body posture (*9*, *10*) recognition, inversion in biological motion is thought to specifically impair holistic processing (*11*), while leaving low-level visual dynamics intact. By disrupting holistic priors we tested a crucial tenet of predictive processing theories that prediction of naturalistic hierarchical input such as biological motion relies on a hierarchy of priors (*20*, *23*, *25*). If instead biological motion prediction relies on general purpose motion extrapolation (e.g., as in (*44–47*)), movie inversion should not affect neural predictions. We hypothesized that inversion specifically attenuates high-level predictions and tested to what extent this attenuation trickles down to lower levels of the hierarchy. Additionally, in a separate manipulation we disrupted kinematics priors of smooth biological motion continuity by temporal piecewise scrambling the videos, which should attenuate prediction across all levels. In other words, attenuation of the predictive representation reveals which priors are actually used for biological motion prediction. These manipulations also allowed us to test an assumption of one specific prediction theory, namely predictive coding, that reducing the predictability of input should result in the representation of more unpredicted information or prediction errors reflecting the actual sensory input (*21*), and which here should be reflected in increased fidelity of post-stimulus representations.

## Results

### Experimental design and behavior

To test the effect of holistic and kinematics priors on predictive neural representations of naturalistic continuous biological motion, we applied dRSA (*35*, *48*) to MEG data of 40 healthy human participants who observed ballet dancing sequences under three different viewing conditions (Fig. 1a). That is, participants observed the biological motion sequences presented either normally (as in (*35*)), up-down inverted, or temporally piecewise scrambled (Fig. 1b). Participants attended the sequences well, as indicated by high performance on occasional catch trials in which the sequence was unexpectedly occluded, after which the participant had to indicate whether the sequence continued correctly (Fig. 1c; top). Importantly, neither in accuracy nor in reaction time was there a significant difference between conditions, indicating that participants paid attention to the biological motion sequence comparably well across the three viewing conditions (Fig. 1d, right; see Supplementary Text for statistical analysis).

**Figure 1.**
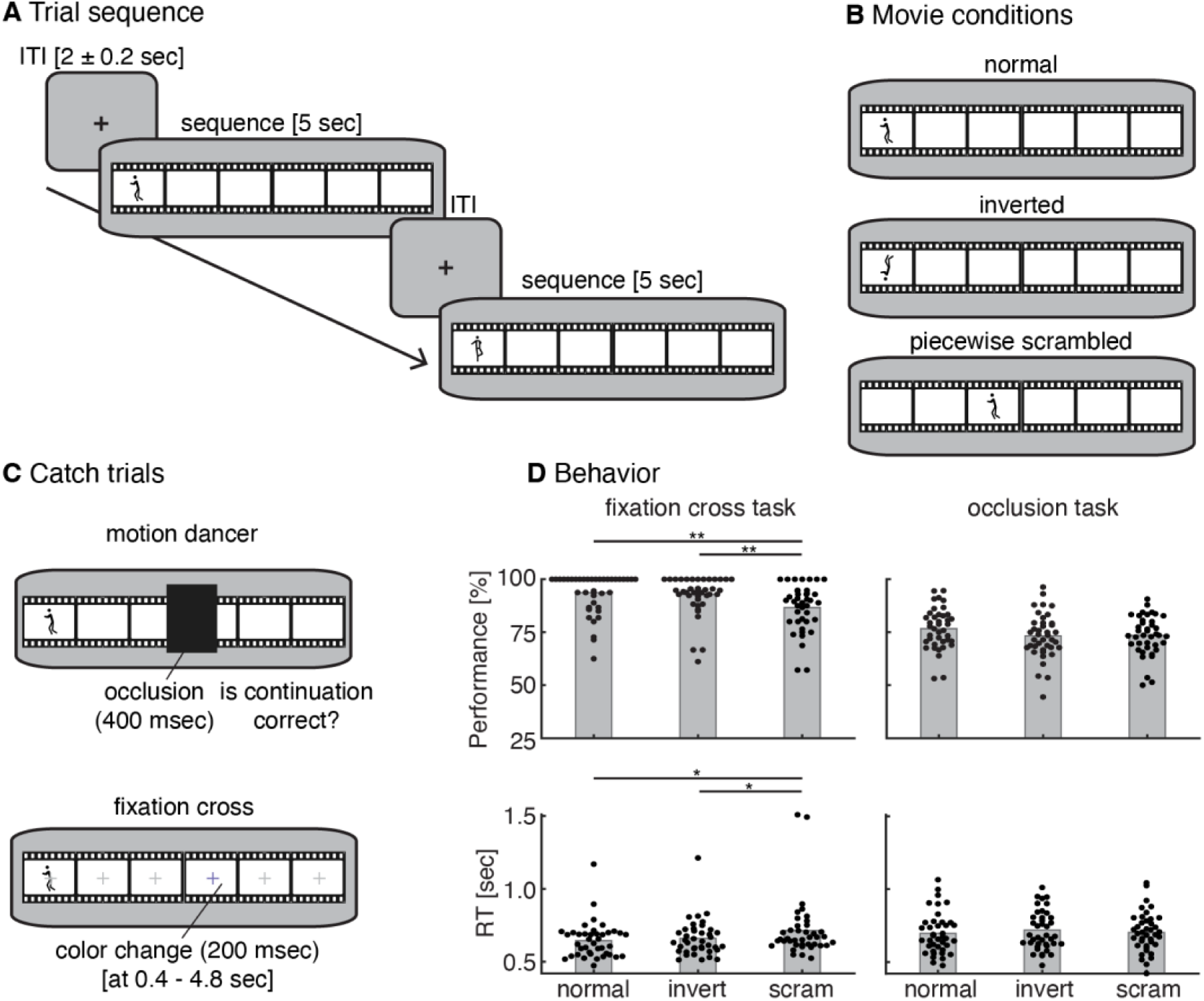
Experimental design and behavior. (**A**) A run consisted of 108 trials alternated by a fixation cross (2 ± 0.2 sec, uniformly distributed). A trial consisted of one of 14 unique 5-sec ballet dancing sequences, such that each unique sequence was shown at least 7 times per run. (**B**) Participants observed the ballet sequences in 3 different viewing conditions (33% of trials per condition); normal, up-down inverted, and piecewise temporally scrambled, with pieces uniformly distributed between 200 – 500 msec. (**C**) Top: To ensure attention to the ballet sequence, in 16 trials per run the video was occluded by a black screen for 400 msec, after which the video continued either correctly, or with an incorrect sequence. Participants (dis)confirmed by button press the correct continuation of the video. Bottom: To encourage fixation, on 8 trials per run the fixation cross changed color for 200 msec at a random time between 0.4 and 4.8 seconds after onset, and participants pressed a button upon detection. (**D**) Behavioral results. Percentage correct and trial-averaged RT on catch trials in top and bottom panels, respectively. RT on the fixation task is counted from color change onset, while RT on the occlusion task is counted from occlusion offset. Dots represent single-participant data (n = 40 healthy human participants). Bars represent the group mean. ITI inter-trial interval. * p < 0.05, ** p < 0.01.

Next, we applied dRSA to the MEG data (for details see Materials and Methods and Fig. S3), to quantify the strength of the match between continuous representations in the brain and continuous feature models of the observed biological motion, across several hierarchical processing levels at the same time. The biological motion videos were modeled at various levels of abstraction across the visual processing hierarchy, from low- level visual features (i.e., pixelwise luminance and motion) to mid- and higher-level, perceptually more invariant features (i.e., 3-dimensional view-dependent and view-invariant body posture and motion), in order to capture a comprehensive characterization of the sequences. dRSA quantifies both the strength of representation and at millisecond-precision how representations temporally relate to (follow or precede) actual events, as reflected in the peak magnitude and peak latency in dRSA latency plots, respectively (Fig. 2 and 4). In case of pure feedforward processing, one expects a lag between the model and the best-matching neural representation (i.e., due to information transfer from retina to V1), as reflected in a peak to the right of the vertical zero-lag midline. Prediction should reduce or even invert (i.e., left of midline) this lag, in which case neural representation predicts the future model state.

**Figure 2.**
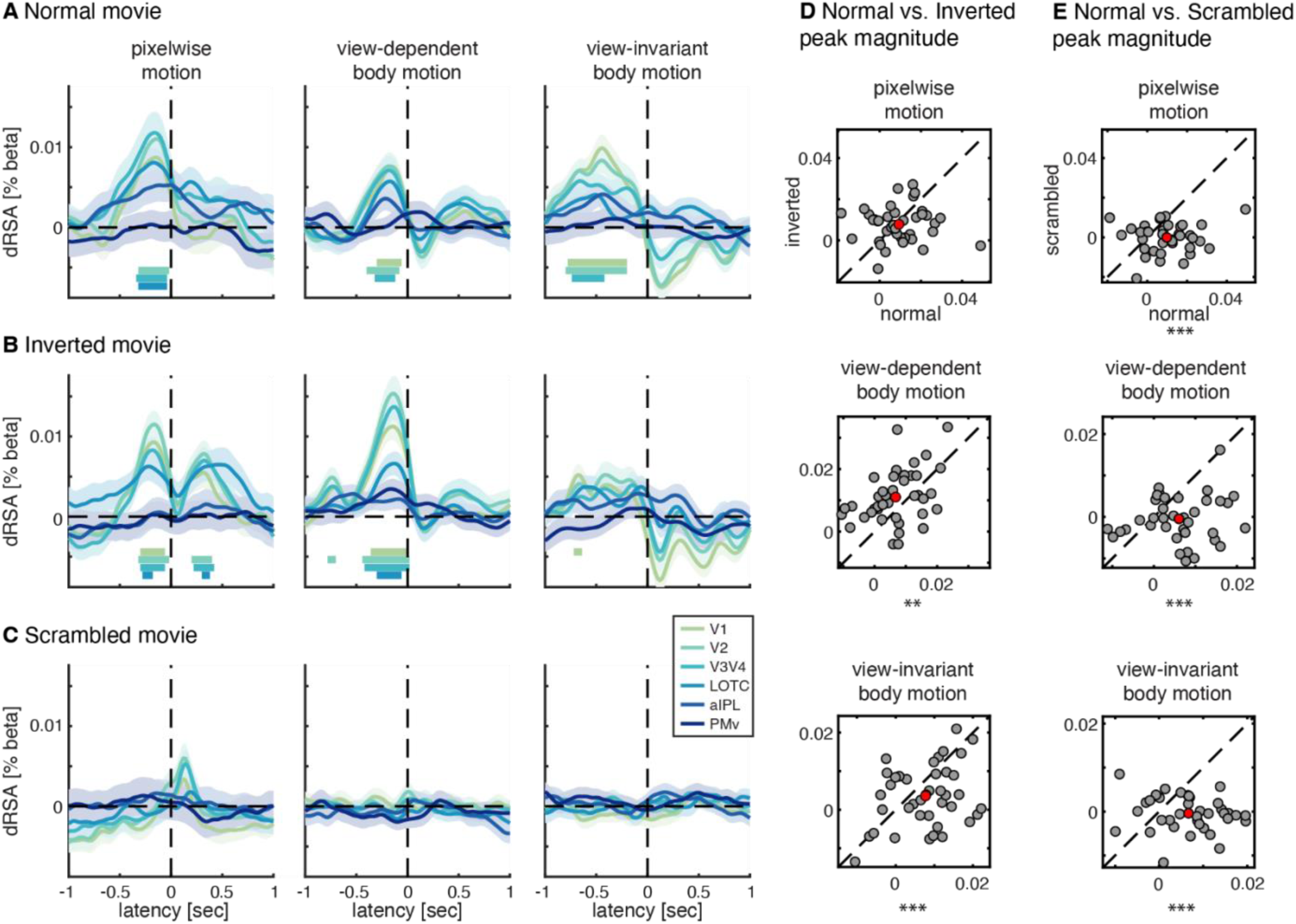
Dynamic RSA results for motion models. Region of interest (ROI)-based analysis, with normalized dRSA regression weights illustrated as latency plots. The three viewing conditions are displayed in rows with normal on top (**A**), inverted in the middle (**B**), and temporally scrambled at the bottom (**C**). Different stimulus models are displayed in columns with pixelwise motion reflecting optical flow vector direction in each pixel, and body motion reflecting 3D motion of 13 kinematic markers placed on the dancer. Lines and shaded areas indicate participant-average and SEM, respectively, with n = 40 in (**A**) and (**B**), and n = 37 in (**C**). Light-to-dark colors indicate posterior-to-anterior ROIs. Horizontal bars indicate beta weights significantly larger than zero (one-sided t-test with p < 0.001 for each time sample), corrected for multiple comparisons across time using cluster-based permutation testing (p < 0.05 for cluster-size), with colors matching the respective ROI line plot. (**D**) Individual-participant (n = 40) peak magnitude in normal versus inverted condition, with peak magnitudes averaged over the first 4 ROIs for pixelwise and view-dependent body motion, and the first 3 ROIs for view-invariant body motion. A dot in the upper left or lower right triangle indicates a larger peak magnitude in the inverted or normal condition, respectively. The red dot reflects the participant average. (**E**) Same as (**D**) but for normal versus scrambled condition (n = 37). ** p < 0.01, *** p < 0.001.

### Biological motion is hierarchically predicted across levels of processing

First, we replicated our previous findings in the normal viewing condition (*35*) and found that the motion of the ballet dancer was predicted at three distinct times that reflected the order in the visual processing hierarchy, such that high-level view-invariant body motion was predicted furthest in advance at ∼500 msec preceding the actual input, while view-dependent body motion and low-level pixelwise motion were predicted at ∼170 and ∼150 msec, respectively (Fig. 2a and Table S1; see Fig. S1 for all models). We ran a [3x4] ANOVA on retrieved jackknifed peak latencies with factors model (pixelwise motion, view dependent and view invariant body motion) and ROI, with the first 4 ROIs (i.e., V1, V2, V3/V4, and LOTC) that showed a significant dRSA effect in at least one of the 3 models. Most importantly, we replicate the previous findings, i.e., a significant main effect of model on the latency of the three predictive peaks (F(1.8,69.5) = 57.98, p < 0.001, ω^2^ = 0.496, BF = 1.375*10^13^).

### Movie inversion attenuates high-level predictions while enhancing mid-level predictions

To test our hypothesis that disrupting holistic priors attenuates biological motion prediction of specifically high-level (view-invariant) body motion, we compared the motion models between the normal and up-down inverted viewing conditions (Fig. 2a and b, respectively). Indeed, we observed a significant reduction in the predictive representation of view-invariant body motion specifically. For statistical comparison we averaged dRSA magnitudes in a 100 msec window surrounding a priori selected predictive peaks based on our previous study (i.e., 110 msec, 180 msec, and 510 msec, for pixelwise motion, view-dependent, and view-invariant body motion, respectively (*35*)), and performed a repeated measures ANOVA for each model, with [2 conditions by 4 ROIs] for pixelwise motion and view-dependent body motion, and [2 conditions by 3 ROIs] for view-invariant body motion, i.e., taking only ROIs where at least one condition showed a significant dRSA effect (Fig. 2a and b; see Fig. 2d for individual-participant peak magnitudes). Most importantly, and in line with our hypothesis, prediction at the highest level of view-invariant body motion was reduced in the inverted compared to normal viewing condition (Fig. 2d; F(1,39) = 16.37, p < 0.001, ω^2^ = 0.095, BF = 5.366). Interestingly, prediction of view-dependent body motion showed the opposite pattern and was stronger for inverted compared to normal (F(1,39) = 8.65, p = 0.005, ω^2^ = 0.054, BF = 7.955). Last, there was anecdotal evidence for no condition difference for pixelwise motion (F(1,39) = 0.41, p = 0.526, ω^2^ < 0.001, BF = 0.349), indicating that low-level motion prediction is minimally, if at all, affected by holistic priors. These effects together suggest that for naturalistic but up-down inverted biological motion, participants still make predictions of (body) motion but rely on the level at which it is possible for them to make predictions (i.e., view-dependent). In other words, disrupting holistic priors specifically attenuates the highest-level predictive representations, while channeling prediction to lower processing levels. Interestingly, this change in neural strategy seems effective, given that both accuracy and reaction time on the occlusion task, which arguably requires prediction, were only numerically lower for inverted than normal (Fig. 1d), with anecdotal-to-moderate evidence for no condition difference.

To test whether up-down inversion also affected the latency of predictive representations, we used bootstrapping to compare the peak latencies (see Methods) between the normal and inverted conditions, with the hypothesis of earlier prediction (more negative latency) for normal compared to inverted movies. Specifically, we tested whether the one-tailed 95% confidence interval of the bootstrapped distribution of normal minus inverted peak latencies was lower than zero. Before bootstrapping, we first averaged the dRSA curves over the same ROIs as before that showed significant effects, since peak latency estimation does not make sense for noisy, non-significant and flat curves. In line with the peak magnitudes, we found that only view-invariant body motion was predicted significantly earlier in the normal compared to inverted condition (−448 vs −352 msec, one-tailed 95% CI: −6 msec), while this was not the case for view-dependent body motion (−165 vs −177 msec, one-tailed 95% CI: 81 msec), nor for pixelwise motion (−217 vs −191 msec, one-tailed 95% CI: 27 msec). Additionally, we computed the centroid latency (i.e., center of mass of the dRSA curve; see Methods), which considers latency differences in representational weight that are not captured by the peak (e.g., for non-symmetrical curves) and is less sensitive to noise than peak latency estimates. Importantly, we qualitatively replicated the results for the centroid latency (see Supplementary Text). Taken together, the latency effects are in line with the magnitude effects, which together show that only view-invariant body motion prediction was affected by inversion.

Besides testing for a representational latency difference between the normal and inverted movies, we could also test whether the neural representational geometry generalized between the conditions, and test whether this shared representation is aligned in time. In order to test this, we ran dRSA directly between the normal and inverted conditions, i.e., by computing the similarity between the neural RDMs of normal and inverted movies across time (Fig. 3a). This analysis captures both the strength of the match between neural representations (peak magnitude), and the latency at which representations matched most strongly, with a peak to the left of the vertical zero-lag midline indicating that the matching neural representation appears earlier in the normal compared to inverted condition. Note that this approach is exactly the same as our main dRSA analysis, except that the model RDM is replaced by the neural RDM of another condition (compare Fig. 3a and Fig. S3). Given that conditions are similar at certain (i.e., orientation-invariant) levels, they should activate similar neural representations for those levels. Additionally, given the earlier neural representation of view-invariant body motion for normal compared to inverted, we hypothesized the shared representation to be earlier in normal compared to inverted biological motion, which would result in a negative peak latency. We used bootstrapping to compare the peak latency against zero in the 100 msec window surrounding zero. Specifically, we tested whether the one-tailed 95% confidence interval of the bootstrapped distribution of peak latencies was lower than zero. Peak latency was increasingly earlier for more anterior areas (Fig. 3b, right panel) with significantly negative latencies at LOTC (−8 msec, one-tailed 95% CI: −1 msec) and V3/V4 (−2 msec, one-tailed 95% CI: −2 msec), but not at V1 (0 msec, one-tailed 95% CI: 12 msec) or V2 (2 msec, one-tailed 95% CI: 9 msec). Since shared variance between the neural RDMs might be partially driven by low-level visual features that are not predicted, we reran the same analysis but used partial correlation as similarity measure to partial out pixelwise luminance and pixelwise motion magnitude, two low-level visual models that showed strong lagged representations (Fig. S1). This analysis revealed qualitatively equivalent results (Fig. 3c), with significant negative latencies at LOTC (−10 msec, one-tailed 95% CI: −1 msec) and V3/V4 (−2 msec, one-tailed 95% CI: −1 msec), but not at V1 (0 msec, one-tailed 95% CI: 8 msec) or V2 (2 msec, one-tailed 95% CI: 7 msec). Importantly, this analysis confirms that neural representations of biological motion sequences at least partly generalize between viewing conditions, and in higher visual areas (V3/4 and LOTC) this shared representation is delayed for the inverted relative to the normal condition. It therefore corroborates the main dRSA results, which suggest that this delay is driven by weaker and later prediction of specifically high-level view-invariant body motion. While this latency difference (up to 10 msec) might seem small compared to the latency difference for view-invariant body motion in the main analysis, this is likely due to the neural-by-neural RDM analysis capturing all shared representation between viewing conditions, including at lower levels at which there are likely no latency differences.

**Figure 3.**
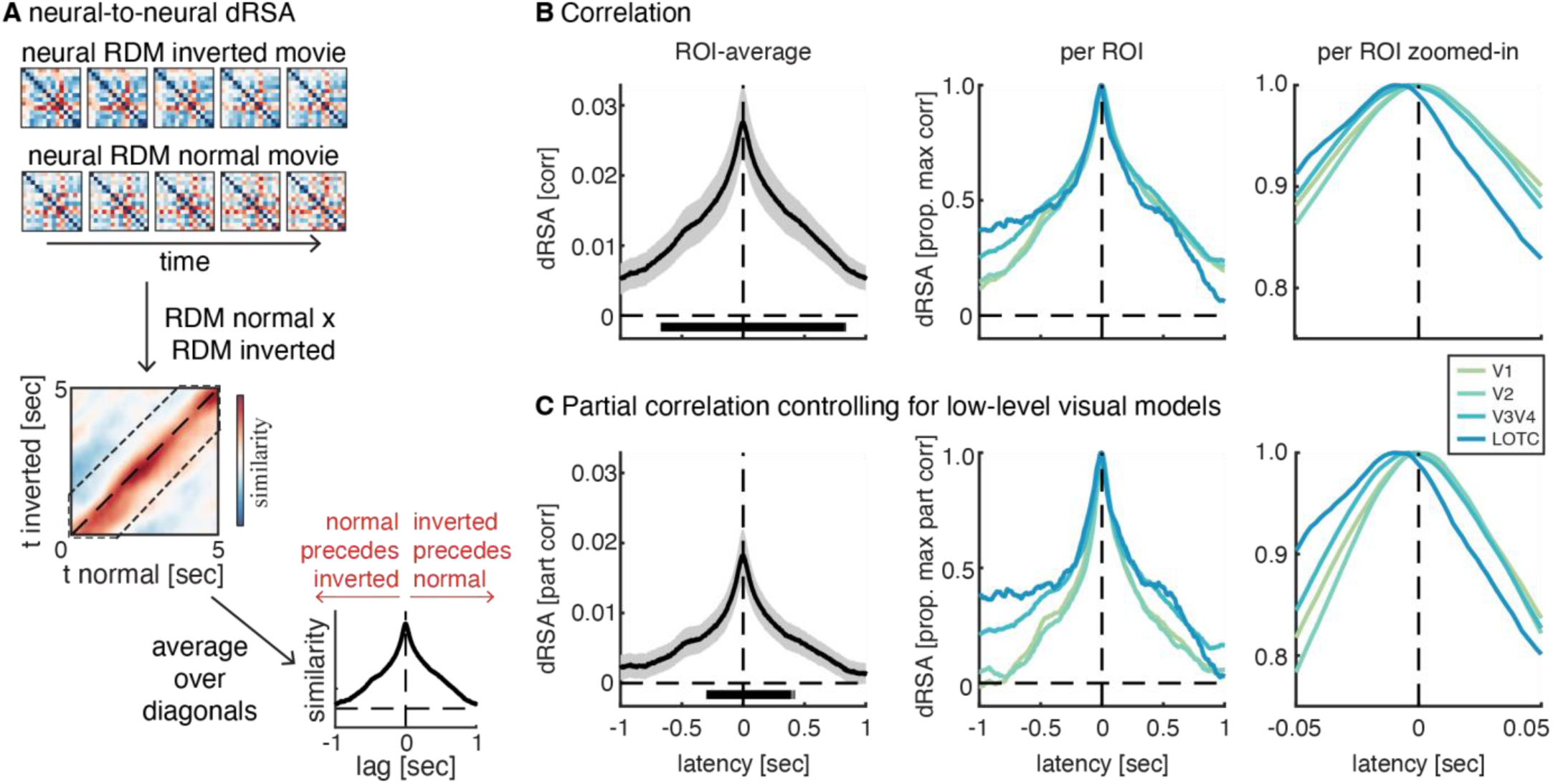
dRSA between neural RDMs of normal and inverted movie conditions. (**A**) Similarity between the neural RDMs of the normal and inverted movie was computed for each normal-by-inverted time point (top), resulting in a normal-time-by-inverted-time dRSA matrix (middle). Last, the 2D dRSA matrix was averaged along the diagonals to create a lag-plot (i.e., lag between normal and inverted RDMs; bottom), in which a peak to the left or right of the vertical zero-lag midline indicates that the neural representation of either normal or inverted is leading, respectively. Similarity between neural RDMs was computed as correlation (**B**), or partial correlation (**C**), where two low-level visual models with lagged representations were partialed out to account for a large part of shared variance between neural RDMs being explained by low-level visual features (i.e., pixelwise luminance and pixelwise motion magnitude; see Methods). In (**B**) and (**C**): The first column reflects the average over the first 4 ROIs. Lines and shaded areas indicate participant-average (n = 40) and SEM, respectively. The horizontal black bars indicate correlation coefficients significantly larger than zero (one-sided t-test with p < 0.001 for each time sample), corrected for multiple comparisons across time using cluster-based permutation testing (p < 0.05 for cluster-size). The second column reflects the participant-average per ROI, with light-to-dark colors reflecting posterior-to-anterior ROIs, while the third column shows the same but zoomed in. Note that the first column reflects absolute correlation coefficients, while the second and third column reflect proportion of maximum correlation coefficient per ROI. Second and third column are smoothed using a 20 msec sliding window for plotting purposes only.

**Figure 4.**
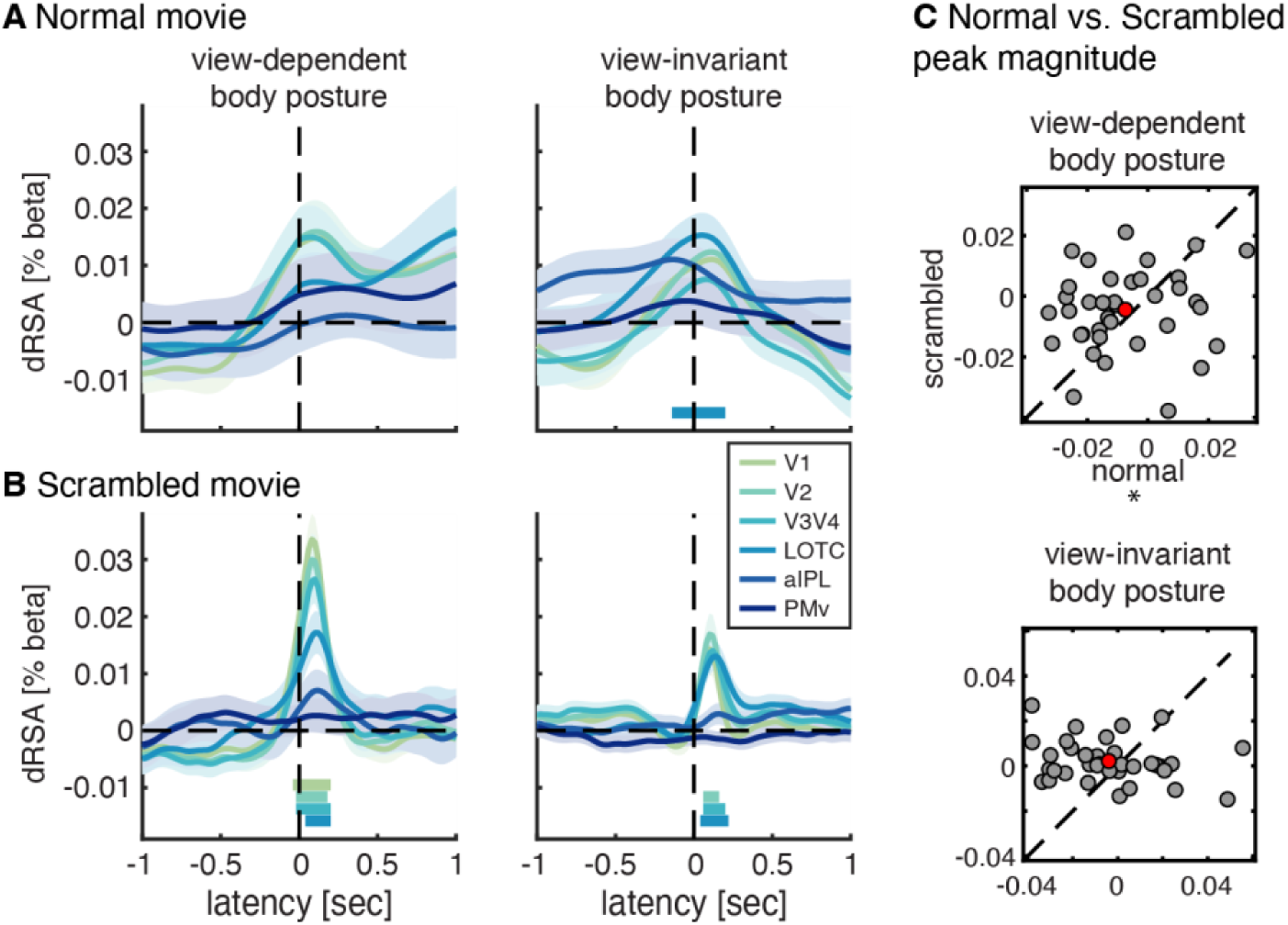
Dynamic RSA results for body posture models. Region of interest (ROI)-based analysis, with normalized dRSA regression weights illustrated as latency plots. Two viewing conditions are displayed in rows with normal on top (**A**) and temporally scrambled below (**B**). Different stimulus models are displayed in columns with body posture capturing 3D position of 13 kinematic markers placed on the dancer. Lines and shaded areas indicate participant-average and SEM, respectively, with n = 40 and 37 in **A** and **C**, respectively. Light-to-dark colors reflect posterior-to-anterior ROIs. Horizontal bars indicate beta weights significantly larger than zero (one-sided t-test with p < 0.001 for each time sample), corrected for multiple comparisons across time using cluster-based permutation testing (p < 0.05 for cluster-size), with colors matching the respective ROI line plot. (**C**) Individual-participant (n = 37) peak magnitude in normal versus scrambled condition, with peak magnitudes averaged over the first 4 ROIs. A dot in the upper left or lower right triangle indicates a larger peak magnitude in the scrambled or normal condition, respectively. The red dot reflects the participant average. * p < 0.05.

### Temporal scrambling shifts representations from predictive to reactive

We hypothesized that if we disrupt kinematics priors by temporal scrambling, all predictive motion representations would be strongly attenuated, which is indeed what we observed (Fig. 2c; i.e., no significant predictive representations). To compare peak magnitudes, we extracted peak magnitudes at a priori selected peak latencies based on our previous study (*35*), for each participant, condition, ROI and model for statistical comparison using a [2 conditions by 4 ROIs by 3 models] repeated measures ANOVA (see Fig. 2e for individual-participant peak magnitudes). We used peak magnitudes instead of average magnitudes in a 100 msec window as for the normal vs inverted comparison, since the normal condition models have broader temporal autocorrelations compared to the scrambled condition (Fig. S2), thus inflating averages. Confirming our hypothesis, magnitudes were indeed significantly attenuated for the predictive motion peaks in the scrambled compared to normal condition (i.e., main effect of condition; F(1,36) = 57.27, p < 0.001, ω^2^ = 0.397, BF = 1.97x10^6^). There was also a minor effect of ROI (F(2,72.2) = 4.98, p = 0.009, ω^2^ = 0.028, BF = 0.633), but moderate evidence for no main effect of model (F(1.5,53.6) = 1.40, p = 0.253, ω^2^ = 0.007, BF = 0.235) and no interactions (p > 0.101, BF < 0.256).

We next tested a key assumption of predictive coding theory, namely that less predictable input should lead to more bottom-up representation of the input. Specifically, we hypothesized that disrupting kinematics priors by temporal scrambling results in an increase in post-event (lagged) representations of static features such as body posture, which is indeed what we observed (Fig. 4). To compare peak magnitudes between the normal and scrambled condition for the body posture models we did not have very reliable a priori peak latencies since these representations were weak in our previous study(*35*). We therefore estimated peak latencies based on the grand-average dRSA latency plots (averaged over participants, normal and scrambled conditions, and the first four ROIs), and extracted peak magnitudes at that latency for each participant, condition, ROI and model for statistical comparison using a [2 conditions by 4 ROIs by 2 models] repeated measures ANOVA (see Fig. 4c for individual-participant peak magnitudes). Confirming our expectation, magnitudes were significantly larger for the lagged posture peaks in the scrambled compared to normal condition (i.e., main effect of condition; F(1,36) = 8.79, p = 0.005, ω^2^ = 0.070, BF = 6.58). There was also anecdotal evidence for a small main effect of model (F(1,36) = 5.67, p = 0.023, ω^2^ = 0.060, BF = 2.47), and for an interaction between model and ROI (F(2.2,80.4) = 3.22, p = 0.040, ω^2^ = 0.022, BF = 1.22), but moderate evidence for no main effect of ROI (F(2.0,71.2) = 2.83, p = 0.066, ω^2^ = 0.017, BF = 0.269) and no other interactions (p > 0.152, BF < 0.459). These results indicate that if the congruent continuation of a biological motion sequence cannot be predicted, the post-event neural representation of (mostly view-dependent) body posture increases. Such post-event representation of unpredicted information (or prediction errors) is in line with predictive coding theory. Last, we tested whether temporal scrambling also affected the latency of body posture representations, using bootstrapping as above to test the expectation that scrambling results in later representations (more positive latency) compared to normal viewing, which would be evidenced by the one-tailed 95% confidence interval of the bootstrapped distribution of normal minus scrambled peak latencies being lower than zero. We found that only the view-invariant body posture was represented delayed in the scrambled compared to normal condition (194 vs 65 msec, 95% CI: −43 msec), while this was not the case for view-dependent body posture (105 vs 133 msec, 95% CI: 101 msec), and we qualitatively replicated this result for the centroid latency (see Supplementary Text).

## Discussion

In this study we tested to which extent the predictive neural representation during biological motion perception depends on holistic and kinematics priors. First, we replicated our previous findings (*35*) that neural representation of normally viewed biological motion is mostly predictive, with time scales matching hierarchical levels of processing such that high-level view-invariant body motion was predicted earliest, while view-dependent body motion and low-level pixelwise motion were predicted increasingly closer to real-time. These findings extend a growing body of evidence for hierarchical prediction (*49–53*), by demonstrating the necessary time scales for such hierarchy. Next, in line with our hypothesis, disrupting holistic priors by up-down inversion attenuated neural prediction selectively at the highest processing level of view-invariant body motion, in terms of both strength and prediction latency, while this was not the case for view-dependent body motion and pixelwise motion. In fact, prediction of view-dependent body motion increased, suggesting that predictions were channeled to this level at which prediction was still possible. Interestingly, this change in neural strategy seemed effective, given that performance on the occlusion task, which arguably requires prediction, was only numerically lower for inverted viewing. Low-level pixelwise motion prediction was not affected by inversion at all, indicating that it does not rely on holistic priors. Next, confirming our hypothesis, disrupting kinematics priors by temporal scrambling completely eliminated neural prediction at all levels. Instead, reactive post-stimulus neural representations for static body posture increased significantly and became more delayed, which likely reflects bottom-up processing of unpredicted sensory input.

Accurate biological motion perception is crucial for our daily social interactions as it helps us understand and predict other’s actions, moods, and intentions (*1*, *2*). While this may be enabled by our priors of other agents and how they move (*4–8*, *11*, *12*, *15–17*, *40*), to date it was unclear whether neural prediction of biological motion relies on such priors, or that general motion extrapolation mechanisms are sufficient. Here we tested the effect of ‘turning off’ our priors. Specifically, we disrupted holistic and gravity priors by up-down inversion, which leaves kinematics intact. Indeed, inversion is an effective way to impede biological motion perception and attenuate its associated neural responses (*6*, *38–41*). Strong inversion effects have also been observed for static body posture (*9*, *10*) and face recognition (*43*, *54*, *55*), which is impaired when inverted, but only when recognition relies on holistic instead of featural processing (*42*), thus evidencing the importance of holistic processing of bodies and faces (*10*, *55*). Interestingly, it seems that for biological motion perception also gravitational priors play an important role. That is, when global configuration is destroyed by spatial scrambling, the inversion effect partly remains due to local biomechanical constraints and acceleration effects (*15*, *38*, *56*). However, as these studies were restricted to motion direction estimation in simple gait, this unlikely generalizes to predictive processing of more complex three-dimensional biological motion. In any case, it is clear that inversion disrupts our holistic priors of other people and how they move. We extend the current literature by showing that specifically the neural prediction of view-invariant biological motion relies on holistic priors, while prediction of view-dependent biological motion and pixelwise motion likely relies more on local motion patterns that are less affected by inversion (*15*, *38*). While this necessity of holistic priors for high-level predictions supports a crucial assumption of hierarchical prediction theories (*20*, *23*, *25*), the intact lower-level predictions suggests that these do not solely rely on a top-down influence of high-level predictions but instead partly originate independently at their respective level of processing.

We tested the effect of kinematics priors on predictive representations by temporal scrambling, and found that all motion prediction, including low-level pixelwise motion, was eliminated. Since pieces were between 200 and 500 msec, short-term kinematics remained intact, and one could expect pixelwise motion prediction (at a latency of only ∼150 msec). It could be that indeed mostly higher-level (view-dependent and -invariant) body motion prediction is attenuated, but that this trickles down to the hierarchically lower pixelwise motion prediction. However, neural prediction of low-level stimuli such as (apparently) moving dots without naturalistic hierarchical embedding is also possible (*44*, *45*). While removing such embedding might attenuate low-level prediction, it seems unlikely to disappear altogether. A more likely explanation is that on average our pieces were too short (i.e., 350 msec) to ‘start up’ any prediction. It is an open question how much time is needed to set up kinematics priors based on local temporal context. Earlier research using temporal scrambling with ∼4 sec pieces in an fMRI movie paradigm found that only higher area responses were disrupted, while responses in early visual areas were not (*57*). However, 4 seconds is too long to investigate low-level visual dynamics. Future research should test how much time is needed to set up our priors, for instance by systematically manipulating piece lengths to see at which length high- and low-level prediction disappears. Interestingly, representational content was dominated by increased and slightly delayed reactive representation for both view-dependent and view-invariant body posture, indicating that if the congruent continuation of a biological motion sequence cannot be predicted, the post-stimulus representation of body posture increases. The here observed stronger post-stimulus representation of unpredicted information supports predictive coding theory, which assumes that sensory input is only processed bottom-up along the visual hierarchy after stimulus occurrence if it has not been predicted, or violates a prediction, i.e., a prediction error (*21*, *24*), which is likely the case here. It is also interesting to note that while here we artificially induce sudden segment boundaries, our daily life is filled with naturalistic event boundaries. Such boundaries are mirrored in boundary-induced activation patterns for short events in sensory regions and long events in high-order areas (*58*), and these boundary-induced activation patterns are similar but advanced on repeated movie viewings (*59*, *60*). Additionally, across compared to within boundaries there is worse behavioral prediction (*61*), delayed eye movements to relevant objects (*62*), and a global neural updating signal (*61*), all indicative of an increase in prediction errors demarcating naturalistic event boundaries. Similarly, observed action events that are isolated from their naturalistic event sequence and randomly scrambled, lead to less top-down activity, and more bottom-up visual responses (*63*). These previous studies thus demonstrate behavioural and neural signatures of reduced prediction and increased prediction errors across (artificial or natural) input boundaries. We extend these findings by unveiling the exact content and latencies of those reduced predictions and increased prediction errors, i.e., eliminated motion prediction and increased and delayed post-stimulus representation of static body posture, respectively.

It is worth mentioning that while we observed reactive representations for body posture in the scrambled condition that we interpret as unpredicted sensory input, a positive latency precludes inferring that a representation solely reflects bottom-up activation, as top-down expectations might very well have modulated its strength (*31*) and processing latency (*64*), i.e., advanced it relative to pure bottom-up processing. It is therefore unclear to what extent these representations are still affected by fast recurrent processing or even prediction for the duration of a video piece. However, a direct comparison with the normal viewing condition does allow us to conclude that post-stimulus representations of body posture are enhanced and delayed, which is indeed in line with predictive coding theory.

To conclude, using a new dynamic RSA approach we demonstrate that high-level view-invariant prediction of biological motion relies on holistic and gravity priors, while in contrast prediction of view-dependent biological motion and low-level visual motion only relies on kinematics priors based on temporally local context. These findings thus provide evidence for a hierarchy of priors, akin to what has been found in other domains (*49–51*), and reveal how these priors shape predictive neural representations of observed biological motion.

## Materials and Methods

### Participants

Forty-three healthy humans (mean age, 28 ± 6 years) participated in the experiment for monetary compensation. Three participants were excluded from analyses due to low behavioral performance (see below). Participants were relatively diverse and consisted of MSc and PhD students, and postdocs, from all faculties of the University of Trento, and originating from a wide range of countries. All but one participant was right-handed, all had (corrected-to-)normal vision and were naïve with respect to the purpose of the study. Sixteen participants identified themselves as male. Sex or gender information was not otherwise collected. As we aimed for our findings to be generalizable, we combined all participants in the group-level statistical analyses. All experimental procedures were performed in accordance with the Declaration of Helsinki and were positively reviewed by the Ethical Committee of the University of Trento. Written informed consent was obtained from all participants. Additionally, written informed consent was obtained from the ballet dancer for the creation and publication of the ballet dancing videos as well as the example frames displayed throughout this article.

### Stimuli

Stimuli consisted of 14 unique 5-sec videos of four smoothly connected ballet dancing figures selected from five unique figures (i.e., passé, pirouette, arabesque, bow and jump; see Table S1 for the figures of each sequence), as used in our previous study (*35*). All ballet figures were presented from several viewpoints to allow separation of view-dependent and view-invariant representations. We recorded 3D positions of 13 infrared markers placed on ankles, knees, hips, shoulders, elbows, wrists, and head of the dancer. See https://github.com/Ingmar-de-Vries/DynamicPredictions for all 14 unique videos and temporally aligned 3D kinematic data.

### Stimulus models

All data analyses were performed in MATLAB (version 2023b; MathWorks). We used stimulus-feature models as described in (*35*). In short, we created a total of 9 stimulus-feature models (Fig. S4), to comprehensively characterize the biological motion sequences at several hierarchical levels of complexity/abstraction from low-level visual information (pixelwise luminance) up to higher-level, perceptually more invariant information (view-invariant body posture and motion). We also included a participant-specific gaze position model to control for eye-movement related variance in the dynamic representational similarity analysis (dRSA). Models based on video data were upsampled from 50 to 100 Hz to match the kinematic data, using shape-preserving piecewise cubic interpolation.

Low-level visual (pixelwise) models (Fig. S4b; left column): RGB values at all 400 × 376 pixels of each video frame were converted to grayscale (i.e., luminance values), smoothed (i.e., spatially low-pass filtered) using a Gaussian kernel with sigma = 5 (i.e., ∼11.75 pixels full width at half max(*36*)), and vectorized, thus resulting in a model vector for each time point. For each time point separately, the dissimilarity between the two vectors of two different videos was computed as 1-Pearson correlation, to generate a single entry in the model RDM (representational dissimilarity matrix) at that time point. This was repeated for each stimulus pair to generate a 14 × 14 RDM at a single time point and subsequently repeated over the 500 time points (i.e., frames) to generate 500 model RDMs (see Fig. S3d for an example). Pixelwise motion was estimated with optical flow vectors (Fig. S4b, middle and right column) using MATLAB’s estimateFlow.m implementation of the Farnebäck algorithm (*65*), which captures the (apparent) motion between subsequent frames at each pixel. Optical flow vectors were spatially smoothed using the same Gaussian kernel as above (*66*), divided in magnitude and 2D direction, and vectorized, before computing the RDM as above.

Kinematic marker models (Fig. S4c): View-dependent body posture (Fig. 4Sc; first column) was operationalized as the 3D positions of the 13 kinematic markers (i.e., 39 features), relative to a coordinate system with origin placed on the floor in the center of the video frame. View-dependent body motion (Fig. 4Sc; third column) was operationalized as the first derivative of view-dependent body posture, i.e., the difference in the 3D marker positions between two subsequent frames. Body motion vectors, therefore, had a magnitude and a 3D direction. View-invariant body posture and motion (Fig. S4c; second and fourth column) were computed by aligning (i.e., minimize sum of squares) the 3D marker structure of one stimulus with the marker structure of the other stimulus, while keeping its internal structure intact, by using translation, and rotation along the vertical axis only, before computing dissimilarity as above. If in two videos the exact same ballet figure is performed from a different perspective, this would result in high dissimilarity of view-dependent body posture, but high similarity of view-invariant body posture. Since dissimilarity is slightly affected by directionality (i.e., dissimilarity(frame1,frame2aligned) ≠ dissimilarity(frame1aligned,frame2)), dissimilarities were computed in both directions and subsequently averaged. View-dependent and view-invariant body acceleration were operationalized as second derivative of body posture. Since these models only explained a minimal amount of variance in the neural data, they are not illustrated in the main results. However, they did correlate with some of the tested models (see subsection Simulations), and we therefore regressed them out when testing each of the other models (see subsection Dynamic representational similarity analysis for details on regression). Note that to prevent artificially large motion vectors (i.e., for pixelwise and body motion) across piece-boundaries in the piecewise scrambled condition, we first computed the models, after which model data was piecewise scrambled to match the stimuli observed by the participants.

Since eye movements are a common covariate in multivariate analysis of neuroimaging data(*67*), even in case of a fixation task (i.e., micro-saccades remain), we included participant-specific RDMs based on gaze-position to ensure that the dRSA results for our models of interest were not due to eye movements. Eye tracker data was downsampled to 100 Hz, missing samples were interpolated (e.g., due to blinks; 1.4 ± 3.0% of all samples; mean ± std across participants), and trials with more than 10% missing samples were rejected (2.7 ± 6.5% of all trials; mean ± std across participants). Finally, trials were averaged per condition over all repetitions of the exact same stimulus. Dissimilarity for the RDM was defined as the Euclidean distance between the four eye position values (i.e., x-, and y-position for the left and right eye), as four values are too low for a reliable correlation estimate. Gaze position did not explain much variance in the neural data (Fig. S1), and simulations confirmed that any remaining variance best explained by gaze position was successfully regressed out when testing each of the other models (see Fig. S2 and subsection Simulations).

### Experimental design and behavioral task

We used Psychophysics Toolbox (version 3) in MATLAB (version 2012b; MathWorks) to create and run the experiment. Trials consisted of 5-sec ballet videos (Fig. 1a) that were separated by blank screens with a white fixation cross presented for 1.8–2.2 s (uniform distribution). Participants kept fixation on a fixation cross displayed on top of the videos while covertly attending the dancer. Videos were presented in three different viewing conditions (33% of trials per condition); normally as in our previous study (*35*), up-down inverted, or temporally piecewise scrambled with pieces randomly and uniformly distributed between 200–500 msec (Fig. 1b). Piece lengths and scrambled indices were different for each participant and each of the 14 videos. They were determined at start of each recording session and stored for offline analysis to allow reconstruction of the exact observed stimuli per participant. For the first three participants we failed to store this information, which thus led to the exclusion of these 3 participants from any analysis involving the scrambled condition. A run lasted ∼14 min and consisted of 108 randomly ordered ballet figures (36 per condition), with the only constraint that the exact same sequence was never directly repeated (i.e., there was always at least one different sequence in between). Attention to the ballet sequence was ensured by means of 16 randomly distributed catch trials per run, in which a black screen occluded the video unexpectedly for 400 msec (Fig. 1c; top). Participants were requested to indicate by button press whether the video continued correctly, or with an incorrect sequence. The occlusion could appear between 2.7 and 4.2 sec after video onset and never crossed the boundary between two ballet figures (which would make the task close to impossible). Additionally, to make the occlusion task possible in the scrambled condition, the scrambled video was always replaced by the normal unscrambled video from 500 msec before occlusion onset. Fixation was encouraged by means of 8 randomly distributed catch trials per run, in which the fixation cross changed color from white to light purple (RGB=[117 112 179]) for 200 msec at a random time between 0.4 and 4.8 seconds after onset, in response to which participants were requested to press a button. The color was chosen such that the change was only visible when fixating, but not from the periphery. Participants received immediate feedback on each catch trial. To ensure continuous attention to the task and fixation, catch trials were uniformly distributed across each run. Participants performed one practice run outside of the MEG scanner, in which 41% of trials were catch trials to ensure enough practice with the task. During MEG, participants performed the behavioral task well, indicating that they attended the dancer and kept fixation (Fig. 1d). Participants performed either 6 or 7 runs depending on attentional fatigue (6.1 on average), which resulted in an average of 659 trials per participant for further analyses (i.e., ∼15.7 trials per unique video per condition). Three participants were excluded from all analyses due to behavioral performance because they responded too fast (< 200 msec) or too slow on the occlusion task in more than 75% of trials (3 participants), or because they had a condition-averaged accuracy on the occlusion task that was lower than the participant-average minus 2 standard deviations (1 participant). This led to a total of 40 remaining participants for the normal and inverted conditions, and 37 for the scrambled condition.

### MEG data collection and preprocessing

MEG was recorded at 1000 Hz using a 306-channel (204 planar gradiometers, 102 magnetometers) VectorView MEG system (Neuromag, Elekta) in a two-layer magnetically shielded room (AK3B, Vacuum Schmelze). A low-pass antialiasing filter at 330 Hz and a DC offset correction was applied online, but no online or offline high-pass filter was applied, since high-pass filtering with too high a cutoff can have unwanted side-effects including temporal spread of multivariate information (*68*). Before MEG recording, we used the Polhemus FASTRAK electromagnetic localization system to digitize the individual-participant head shape, i.e., the position of three anatomic landmarks (nasion, left and right preauricular points), five head position indicator coils and at least 350 additional points uniformly spread across the participant’s head. We digitized landmarks and coils twice to minimize error. To allow offline MEG – MRI co-registration, we acquired the head position at start of each run by passing a small current through the coils. We collected binocular gaze position at 1000 Hz using the SR-Research Eyelink Plus eye tracker. Stimuli were presented using a Vpixx PROPixx projector. All hardware was connected to a DataPixx input-output hub (Vpixx Technologies) to present stimuli, and collect data and button presses with minimal delays, and to store stimulus-onset triggers together with the MEG data. Additionally, a photodiode in the top left corner of the screen (i.e., outside of the video frame and invisible to the participant), registered a square on the screen underneath the photodiode changing from black to white at the onset of each single video frame, for realigning MEG signals to these photodiode signals offline.

As a first preprocessing step, bad sensors (e.g., noisy or with many SQUID jumps) were automatically detected using MNE python’s find_bad_channels_maxwell algorithm (*69*), to detect a maximum of 12 noisy sensors for interpolation. Neuromag’s MaxFilter implementation (version 2.2) of Signal Source Separation (SSS (*70*)) was used to remove external noise, interpolate bad sensors, and spatially align the head position inside the helmet across runs. All further preprocessing was done using Brainstorm (version 3)(*71*) and Fieldtrip (version 20191113)(*72*) in MATLAB (version 2023b; MathWorks), as well as custom-written MATLAB scripts (shared at https://github.com/Ingmar-de-Vries/HierarchicalPriors). MEG data were loaded into Brainstorm, after which spatial co-registration with participant-specific anatomical MRI scans was refined using the 350+ digitized head points, data were filtered for line noise (50 and 100 Hz, 0.5 Hz filter bandwidth), down-sampled to 500 Hz, and cleaned using independent component analysis (ICA). ICA was applied separately to magneto-and gradiometers, and on a temporary data version that was downsampled to 250 Hz and band-pass filtered at 1–100 Hz to speed up computation and improve ICA, respectively. An average of 2.2 blink and eye-movement, 1.5 cardiac and 0.3 noise related components were detected by visual inspection and removed from the original data. Next, segments were automatically marked as bad using Brainstorm based on low-frequency noise (i.e., 1–7 Hz; e.g., movement or SQUID jumps) or high-frequency noise (i.e., 40–240 Hz; e.g., muscle), with the sensitivity parameter set to 4 or 5 depending on visual inspection of detected segments (participant-average of 4.8 for each frequency band). Next, continuous data were epoched (−1.5 to 6.5 sec locked to video onset), single-trial baseline corrected (−500 to 0 msec), and previously detected bad segment were converted to NaN. This resulted in an average of 614 out of 659 completely clean trials. We then applied source reconstruction on these clean trials only (see below) and only stored the inversion kernels (i.e., 306 x 15.000 matrix) for efficiency. Last, sensor-level data were exported in Fieldtrip format and Fieldtrip functions were used to realign epochs to photodiode onset, temporally smooth data using a 18 msec boxcar kernel, downsample to 50 Hz, and finally average over all ∼15.7 repetitions of the same unique video per condition. Note that for the average we ignored the bad segments (i.e., NaNs), as well as the time window after catch-trial onset in the catch-trials.

### Source reconstruction and ROI selection

We used minimum-norm estimation (MNE) in Brainstorm with default settings (*73*), to estimate 15,000 source signals distributed on the cortical surface. For 30 participants we used participant-specific anatomical T1 MRI scans, and if unavailable the ICBM152 standard brain for the remaining 10 participants, to construct 3D forward models. Scans were obtained using a Siemens Prisma 3T. Fiducial points were marked automatically in MNI space, after which 3D brains were reconstructed using CAT12 version 2170 (*74*). For the 10 participants without individual scan, the standard brain was warped to the participant’s head shape as estimated from the +350 digitized head points. Overlapping spheres were used as forward model. Noise covariance matrices were computed from the baseline-normalized single trials using the −1000 to 0 msec pre-video time window and using both gradio- and magnetometers. Non-normalized current density maps were computed as measure of source activity, with source direction constrained to be normal to the cortex, thus resulting in a single signal at each source. Next, the 15,000 sources were parcellated into 360 parcels according to the Human Connectome Project (HCP) atlas (*75*). Six regions of interest (ROIs) that consisted of one or more parcels were selected a priori consistent with our previous study (Fig. S5) (*35*), i.e., three low-level visual areas (i.e., V1, V2 and V3+4), and three areas from the action observation network, namely the lateral occipitotemporal cortex (LOTC), the anterior inferior parietal lobe (aIPL), and the ventral premotor cortex (PMv), because this network is involved in processing observed actions (*76–78*). ROIs were comparable in size with on average 484 sources (min–max = 403–565). Source signals were used as features to compute a neural RDM for each ROI, using 1–Pearson correlation as dissimilarity after centering the data.

### Dynamic representational similarity analysis

We applied the same dRSA pipeline with similar parameters as validated in our previous study (*35*), but describe it here shortly for completeness. Time-resolved neural and model RDMs were computed as described above (see also Fig. S3d), resulting in RDMs of 91 features (i.e., 14 × 14 stimuli) by 250 time points (at 50 Hz). RDMs were temporally smoothed using a 30 msec boxcar kernel to improve SNR, and subsequently centered per time point, and standardized across all time points at once to equalize scales between RDMs while keeping temporal structure inherent to a certain model intact. Next, the similarity between neural and model RDMs was computed at each neural-by-model time point (Fig. S3e; bottom panel), and averaged over time for each neural-to-model time lag (i.e., for each diagonal in the 2D dRSA matrix) within the range of −1 to 1 sec. This resulted in a latency-plot (Fig. S3e; top panel), in which a peak to the right of the vertical zero-lag midline indicates that the neural RDM is most similar to a model RDM earlier in time, whereas a peak on the left indicates that the neural RDM is predicting a future model RDM. The different stimulus models inevitably share variance at any given time point, but also across time points due to temporal autocorrelation within models (see subsection Simulations and Fig. S2a and c), which are not problems with dRSA per se, but rather inherent to naturalistic continuous stimuli. We therefore used principal component regression (PCR) to compute similarity, which estimates variance in the neural RDM best explained by each separate model RDM (Fig. S2b and d), while minimizing variance better explained by other models. The neural RDM at a given time point t_y_ is the response variable Y (91 features by 1 time point), while the regressors consist of the model RDM of interest at t_x_ (91 x 1), the other 9 model RDMs across latencies from −1 to 1 sec (91 x N_time_ per model), plus the model RDM of interest itself at distant time points (i.e., minimum distance from t_x_ at which the model shared less than 10% variance with itself at t_x_) to minimize effects of model autocorrelation. The regression weight for the model RDM of interest at t_x_ is the measure of similarity with the neural RDM at t_y_. Since this results in >900 regressors (i.e., 91 x 900), we used principal component analysis (PCA) to reduce dimensionality. First, we ran PCA separately per model and selected those components capturing at least 0.1% of total variance. We then combined models and ran a second PCA from which only the first 75 components were selected (to stay well below the theoretical limit of 91 features). These 75 components were used as predictor variables in a linear least-squares regression, after which the PCA loadings were used to project the component regression weights back onto the original regressors, in order to extract the regression weight for the model RDM of interest at t_x_. PCR strongly reduces dimensionality of the regressors while maintaining almost all variance. Additionally, it decorrelates regressors which prevents multicollinearity. Crucially, simulations confirmed that PCR was effective at estimating a specific model representation while minimizing variance better explained by other models (see subsection Simulations and Fig. S2).

### Temporal subsampling

To increase SNR and generalizability of results we included an additional important step in the dRSA pipeline. At any given time point stimulus-specific feature trajectories determine how far in advance a stimulus can be predicted, which differs for each of the 14 videos. The RDM structure at any time point therefore depends on the exact arbitrary pairwise alignment of the ballet sequences, which in turn causes the pattern across stimulus time (i.e., along the diagonal) in the 2D dRSA matrix to be heterogeneous and idiosyncratic. Since we’re not interested in these specific 14 stimuli and their arbitrary alignment per se, we added a temporal subsampling and realignment step to mimic an infinite number of stimuli and thus minimize the idiosyncrasy of our dRSA results (Fig. S6). Specifically, across 1000 iterations a random 3-sec segment was extracted from each video (Fig. S6a; orange boxes). While a different time window was selected per stimulus, the same time window was used for neural and model data within a stimulus, thus keeping their temporal alignment intact (Fig. S6a; dotted vertical orange lines). Next, the new 14 3-sec segments were realigned, and RDMs were computed at each realigned time point t_R_ (Fig. S6b and c), after which similarity between neural and model RDMs was computed for each realigned neural by model time point as described above. This procedure results in a different pairwise alignment of the 14 stimuli on each iteration, thus creating different idiosyncratic dRSA latency curves on each iteration with different shapes and peak latencies (see Fig. 7d in De Vries et al. 2023 (*35*) for an illustration). In the last step dRSA latency curves were averaged across the 1000 iterations, thus reducing dependency of results on arbitrary pairwise alignments of biological motion sequences, make the results more generalizable, and increase SNR. Note that this whole procedure was done separately per participant.

### Statistical analysis

First, we tested for which ROI–model–condition combination the dRSA latency curve (i.e., beta weights) were significantly higher than zero. We performed group-level non-parametric permutation testing (25,000 permutations per analysis) with cluster-based correction for multiple comparisons (*79*), with t-values from a one-tailed t-test as test statistic and the sum over t-values within a cluster as measure of cluster size. Clusters were made up of connected time points with p-values below 0.001, and significance of cluster-size was tested at alpha = 0.05. Only significant clusters larger than what might be expected by chance survive this procedure. Significant intervals in the dRSA latency plots are indicated by think horizontal bars with colors matching the respective ROIs (Fig. 2, 4 and S1).

The two main measures of interest we used to compare conditions are the magnitude and latency of the peaks, which capture the strength of the representation, and the timing relative to the stimulus event (i.e., how lagged or predictive) at which the representation is strongest, respectively. Given that peak latencies are difficult if not impossible to capture for weak representations (i.e., a nonsignificant dRSA effects), we restricted the condition comparisons to ROIs where at least one condition showed a significant dRSA effect (Fig. S1). To compare magnitudes of predictive peaks between the normal and inverted conditions, we first averaged dRSA values in a 100 msec window surrounding a priori selected predictive peaks based on our previous results with the same stimuli (*35*) (i.e., ∼110 msec, ∼180 msec, and ∼510 msec, for pixelwise motion, view-dependent, and view-invariant body motion, respectively), and performed a repeated measures ANOVA for each model separately, since time-averages are not comparable between models with different temporal autocorrelation profiles, i.e., averages are inflated in case of broader temporal autocorrelation (Fig. S2). This resulted in a [2 condition by 4 ROIs] ANOVA for pixelwise motion and view dependent body motion, and a [2 conditions by 3 ROIs] ANOVA for view invariant body motion. To compare peak magnitudes between the normal and scrambled condition for the body posture models we did not have very reliable a priori peak latencies since these representations were weak in our previous study. We therefore estimated peak latencies based on the grand-average dRSA latency plots (averaged over participants, normal and scrambled conditions, and the first four ROIs), and extracted peak magnitudes at that latency for each participant, condition, ROI and model for statistical comparison using a [2 conditions by 4 ROIs by 3 models] repeated measures ANOVA for predictive peaks and [2 conditions by 4 ROIs by 2 models] ANOVA for lagged body posture peaks. ANOVAs were run in JASP version 0.19.3 with default parameter settings (e.g., Cauchy prior width set to 0.707) (*80*). Frequentist ANOVAs were used for hypothesis testing, while the strength of evidence in favor of or against a given effect was further estimated as Bayes Factor (BF) using Bayesian ANOVAs. Violations of the sphericity assumption were corrected using Huynh-Feldt. For Bayesian ANOVAs we analyzed the effects for matched models, i.e., a direct comparison of the evidence between all models that include a given factor and all equivalent models without that factor.

We used bootstrapping to compare peak latencies between conditions (or against zero for the direct comparison between neural RDMs of normal vs inverted; Fig. 3), a common approach that is sensitive to capture small latency differences (e.g., (*37*)). Specifically, across 10,000 iterations we extracted a random sample (with replacement) of size N from our N participants, estimated the peak latency difference between conditions for each bootstrapped participant, and averaged over participants. These 10,000 bootstrapped averaged peak latency differences were then used to create a distribution, and to test whether the 95% confidence interval of the bootstrapped distribution included zero. Additionally, we performed the exact same bootstrapping analysis on the centroid latency, which is the center of mass of the dRSA curve, or the latency at which the area under the curve to the right and left side of this latency is equal. The centroid latency might be less sensitive to noise than peak latency estimates, as it considers representational weight that is not captured by the peak (e.g., for non-symmetrical curves). Last, for an exact replication of our previous results regarding hierarchical time scales of predictions in the normal condition only (*35*), we computed retrieved jackknifed peak latencies and ran a [3 conditions by 4 ROIs] repeated measures ANOVA.

### Simulations

While we used simulations in our previous study to validate the regression-based dRSA pipeline (*35*), we computed them here again for completeness, now including the new conditions. Most importantly, the simulations demonstrate that principal component regression (PCR; see subsection Dynamic representational similarity analysis) is effective at estimating variance in the (simulated) neural RDM best explained by each separate model RDM, while minimizing variance better explained by other models (compare Fig. S2a and c with b and d, respectively). Specifically, each simulated neural RDM consisted of a single model RDM at zero lag with zero added noise. We ran dRSA with PCR beta weights as similarity measure (Fig. S2b and d), as well as correlation for comparison (Fig. S2a and c). Note the nonzero off-midline correlations both across and within models, which demonstrate the complex patterns of shared variance across time, which in turn limits unambiguous inference if correlation is used. Instead, PCR is effective at extracting only the model of interest from the simulations, while largely regressing out other models. Note that the models for the normal and inverted condition are identical, while the scrambled condition inevitably results in models with narrower temporal autocorrelation. Interestingly, these simulations indicate a different maximum possible PCR beta weight for each model and for the normal/inverted vs scrambled condition (i.e., see zero-lag peak magnitudes in Fig. S2b and d). We used these maximum possible beta weights to normalize the observed beta weights against (i.e., % max beta in Fig. 2, 4 and S1), which allows better comparison between models or conditions with different maximum possible regression weights.

## Acknowledgments

We thank Natasha Bertelsen who volunteered to help with data collection, allowing us to finish just before planned summer closure of the MEG scanner. We thank Carlo Miniussi, former director of CIMeC, where the experiment was conducted, for agreeing to keep the MEG scanner operational 2 weeks longer than planned before summer closure. We thank Danielle E. Parrott for kindly serving as the ballet dancer in our video stimuli.

## Funding

European Commission Marie Skłodowska-Curie Actions postdoctoral fellowship, PredictiveBrain, 101060807 (IEJV).

European Research Council, SURPRISE, 101000942 (FPL).

Netherlands Organisation for Scientific Research (NWO), Vici, VI.C.231.043 (FPL).

Fondo Italiano per la Scienza, DYNAMO, FIS00003032 (MFW).

## Author contributions

Conceptualization: IEJV, FPL, MFW

Methodology: IEJV, FPL, MFW

Investigation: IEJV

Visualization: IEJV

Supervision: FPL, MFW

Writing—original draft: IEJV

Writing—review & editing: IEJV, FPL, MFW

## Competing interests

The authors declare that they have no competing interests.

## Data and materials availability

The experimental code and stimuli, all MEG, eye-tracking, and behavioral data, and all analysis code needed to run the presented analyses and evaluate the conclusions are freely available. Code is available at: https://github.com/Ingmar-de-Vries/HierarchicalPriors. Stimuli are available at https://github.com/Ingmar-de-Vries/DynamicPredictions. The data will be shared in BIDS format on the Radboud Data Repository (https://data.ru.nl/).

## Supplementary Materials for

### Supplementary Text

#### Behavioral results

Behavioral results on the catch trials are illustrated in Fig. 1d in the main text. On the occlusion task, participants responded 77 ± 10, 73 ± 11 and 74 ± 9 percent correct (mean ± std) on the normal, inverted and scrambled movie conditions, respectively (Fig. 1d; top, right), with anecdotal evidence for no difference between conditions (F(2,78) = 2.05, *p* = 0.136, ω^2^ = 0.014, BF = 0.467). Similarly, there was moderate evidence that participants responded equally fast on all three conditions in the occlusion task (F(2,78) = 1.68, *p* = 0.194, ω^2^ = 0.001, BF = 0.313), with reaction times of 697 ± 141, 716 ± 133, and 706 ± 135 msec on the normal, inverted, and scrambled conditions, respectively (Fig. 1d; bottom, right). On the fixation task, participants responded 100 ± 12, 94 ± 8, and 89 ± 14 percent correct (median ± IQR) on the normal, inverted and scrambled conditions, respectively (Fig. 1d; top, left), with strong evidence for a difference between conditions (F(2,78) = 7.79, *p* < 0.001, ω^2^ = 0.074, BF = 44.39), which a post-hoc test indicated was driven by a lower performance on the scrambled compared to both the normal condition (t(39) = 3.54, *p* = 0.003, d = 0.686, BF = 29.16) and the inverted condition (t(39) = 3.08, *p* = 0.008, d = 0.583, BF = 9.35). Similarly, there was strong evidence for a difference in reaction time between conditions in the fixation task (F(2,78) = 7.01, *p* = 0.002, ω^2^ = 0.030, BF = 19.34), with reaction times of 640 ± 145, 736 ± 126, and 757 ± 108 msec on the normal, inverted, and scrambled conditions, respectively (Fig. 1d; bottom, left). This difference was again driven by a slower reaction time on the scrambled compared to both the normal condition (t(39) = 3.10, *p* = 0.011, d = 0.441, BF = 9.97), and the inverted condition (t(39) = 2.44, *p* = 0.039, d = 0.355, BF = 2.31). The reduced performance on the fixation task for the scrambled condition is likely due to the color change of the cross being harder to detect amidst the sudden frame changes between scrambled video pieces. Importantly, the lack of a difference in performance on the occlusion task indicates that participants paid attention to the ballet dancer equally well across the three viewing conditions. Last, participants did not systematically choose to either focus only on the fixation cross, or only on the ballet dancer, which would be reflected in a negative correlation between tasks. Instead, there was moderate evidence for no correlation for percentage correct (r = −0.008, *p* = 0.959, BF = 0.197), and a positive correlation for reaction time (r = 0.440, *p* = 0.005, BF = 9.65), which suggests that participants divided attention across tasks as intended.

#### Centroid latency

Our main estimate of representational latency was based on the latency of the peak in our dRSA latency plots (Fig. 2 and 4). However, peak latency estimation is inaccurate, sensitive to noise, and might not capture the weight of the representational timing in case of non-symmetrical curves. We therefore also computed the centroid latency (i.e., center of mass of the dRSA curve; see Methods), which considers latency differences in representational weight that are not captured by the peak. Importantly, for all our peak estimations the centroid analysis showed qualitatively the same pattern. That is, only view-invariant body motion was predicted significantly earlier in the normal compared to inverted condition (−498 vs −418 msec, 95% CI: −16 msec), while this was not the case for view-dependent body motion (−174 vs −172 msec, 95% CI: 38 msec), or for pixelwise motion (−239 vs −227 msec, 95% CI: 16 msec). For the body posture models only view-invariant body posture was delayed in the scrambled compared to normal condition (176 vs 93 msec, 95% CI: −20 msec), while this was not the case for view-dependent body posture (94 vs 187 msec, 95% CI: 149 msec).

**Fig. S1.**
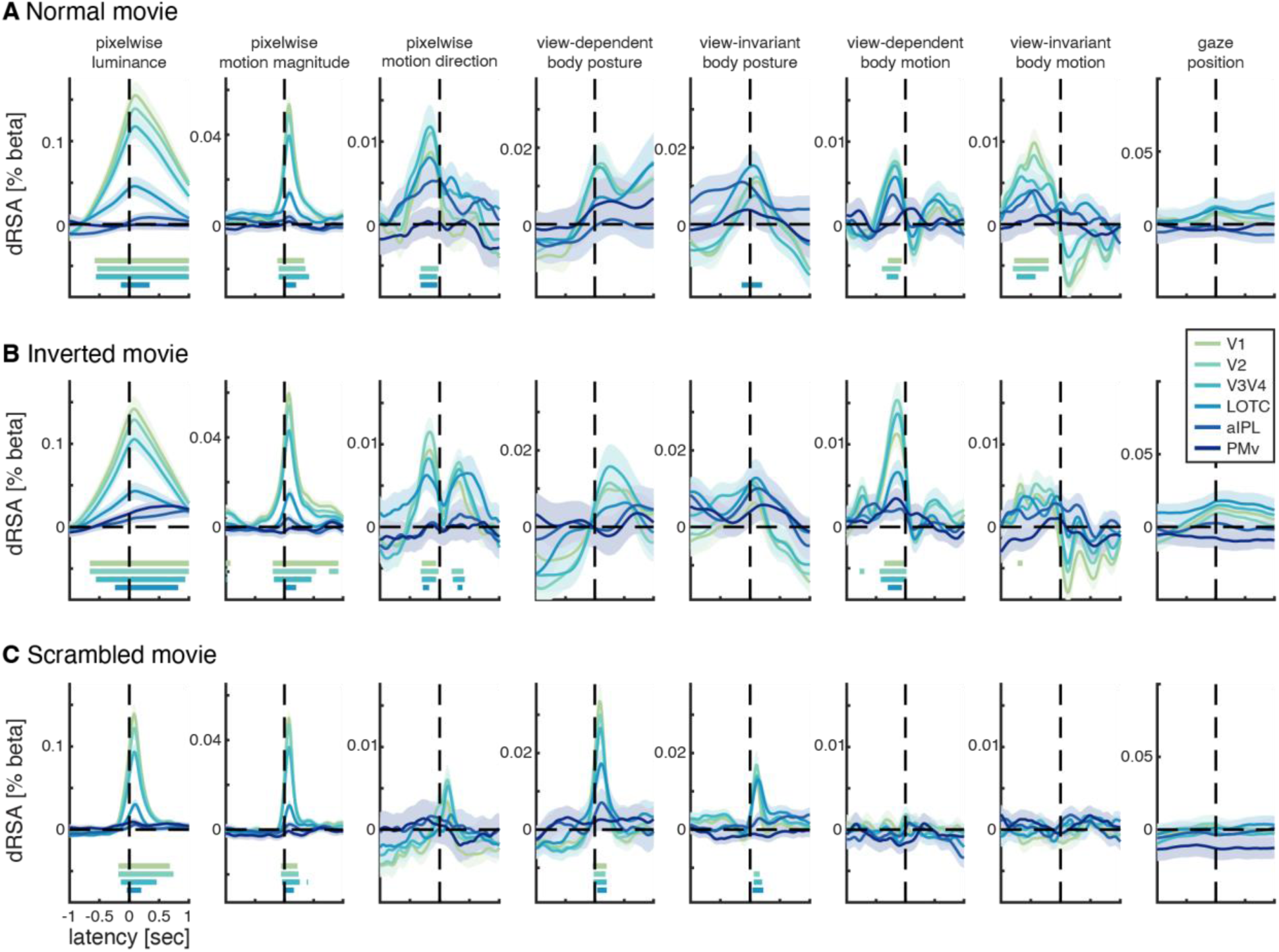
dRSA results for all models. Region of interest (ROI) analysis, with normalized dRSA regression weights illustrated as latency plots for all tested models. The three viewing conditions are displayed in rows with normal on top (**A**), inverted in the middle (**B**), and temporal piecewise scrambled at the bottom (**C**). Different stimulus models are displayed in columns. Lines and shaded areas indicate participant-average and SEM, respectively, with n = 40 in (**A**) and (**B**), and n = 37 in (**C**). Light-to-dark colors indicate posterior-to-anterior ROIs. Horizontal bars indicate beta weights significantly larger than zero (one-sided t-test with p < 0.001 for each time sample), corrected for multiple comparisons across time using cluster-based permutation testing (p < 0.05 for cluster-size), with colors matching the respective ROI line plot.

**Fig. S2.**
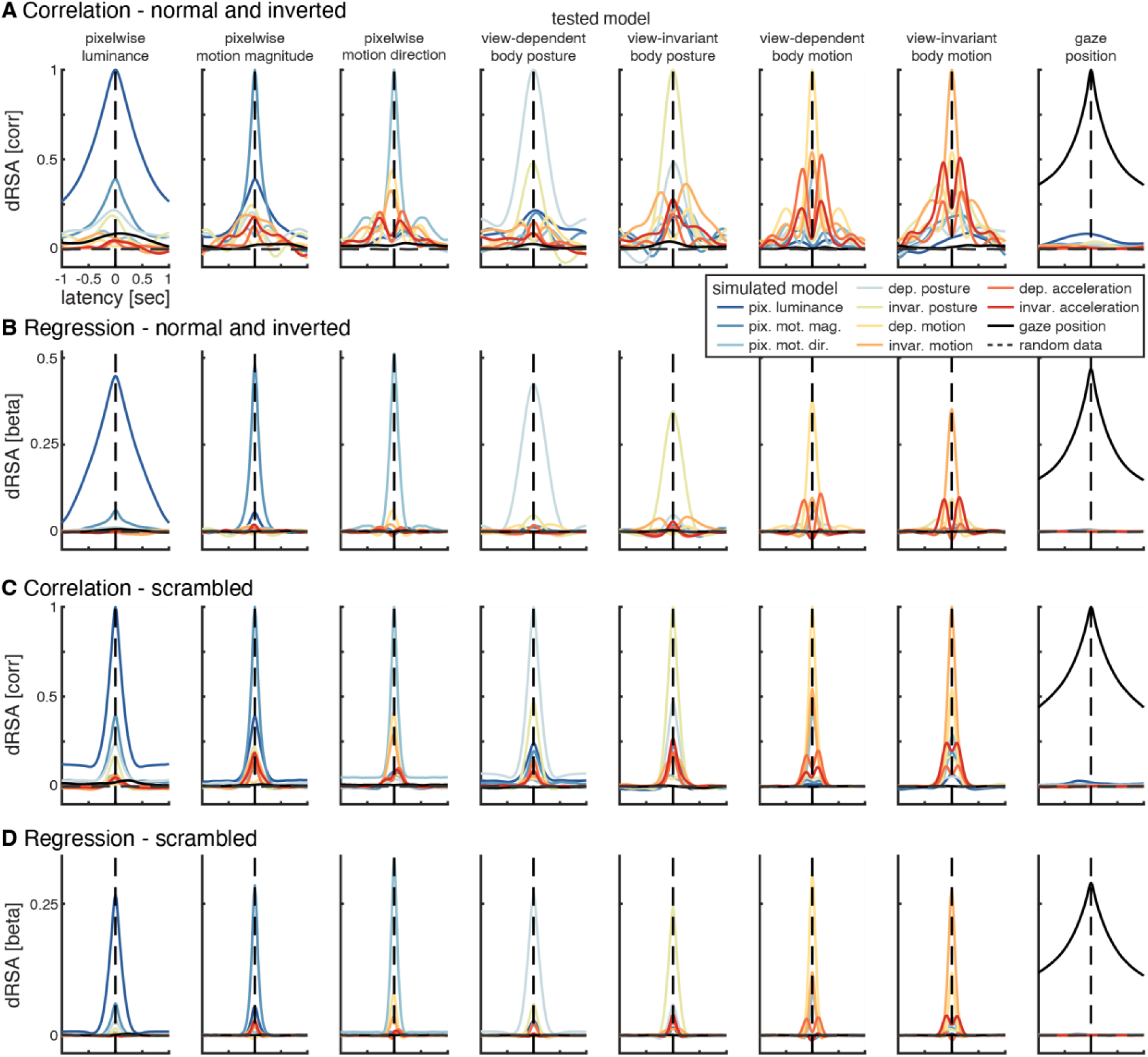
Simulations. (**A**) dRSA on simulated data using simple correlation as similarity measure, in the normal and inverted viewing conditions. Note that the model RDMs do not differ between the normal and inverted viewing conditions. Colors indicate separate simulations of individual model RDMs. Columns indicate which model was tested. These results illustrate the effect of shared variance between various models and the effect of temporal autocorrelation within a given model. (**B**) Same but with regression weight as similarity measure for dRSA using principal component regression (PCR; see “Materials and Methods” for details). In short, simulated neural RDMs are exactly the same as in (**A**), but at the final step in dRSA (i.e., comparing neural and model RDMs) all models are included in a single regression, and only the beta weight of the tested model (columns) is illustrated. (**C**) and (**D**) Same as (**A**) and (**B**), respectively, but for the piecewise scrambled viewing condition. Importantly, these results indicate that PCR is effective at extracting only the model of interest from the simulated neural RDM, while largely regressing out the other models.

**Fig. S3.**
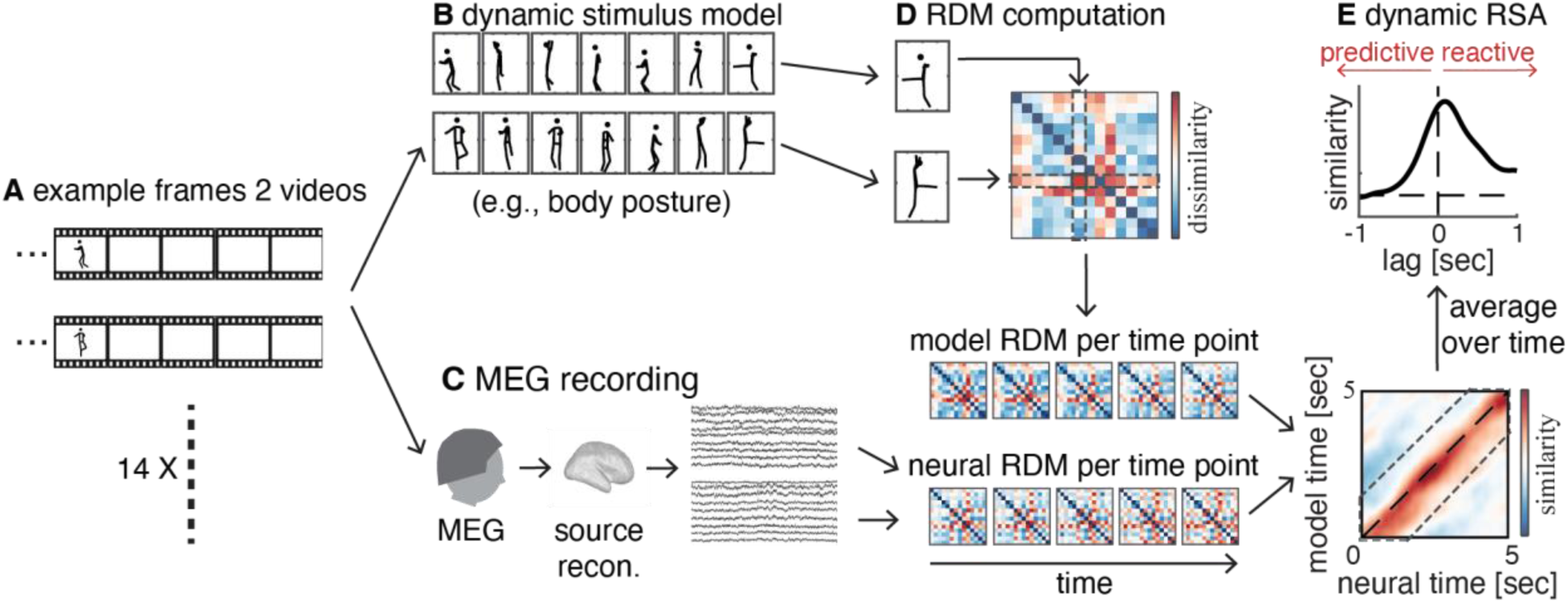
Dynamic representational similarity analysis (dRSA) approach. Figure adapted from De Vries et al. 2023, Nat Commun, CC BY 4.0 (1). (**A**) Subjects observed ∼46 repetitions of 14 unique 5-sec dancing videos (Table S1) in 3 different viewing conditions (Fig. 1a and b) during MEG (i.e., ∼15 repetitions per video per condition). (**B**) Stimuli were characterized at different levels using dynamic stimulus models (e.g., body posture; see Fig. S4 for all models). (**C**) Individual-subject source-reconstructed MEG signals within regions of interest (ROI; see Fig. S5 for ROI definitions) were used as features for subsequent steps. (**D**) Neural and model representational dissimilarity matrices (RDMs) were created at each time point based on pairwise dissimilarity in neural responses to the 14 stimuli and pairwise dissimilarity in stimulus feature models, respectively. Bottom: model and neural RDMs are shown for 5-time points. (**E**) Similarity between neural and model RDMs was computed for each neural-by-model time point (lower panel), using regression weights to test a specific model RDM, while regressing out other covarying model RDMs. This approach was validated through simulations (see subsection ‘Simulations’ and Fig. S2). Last, the 2-dimensional dRSA matrix was averaged along the diagonal to create a latency-plot (i.e., the lag between neural and model RDM; upper panel), in which peaks to the right or left of the vertical zero-lag midline reflect reactive or predictive neural representations, respectively. These dRSA latency plots are computed separately for each subject, ROI, model and condition, and then statistically tested.

**Fig. S4.**
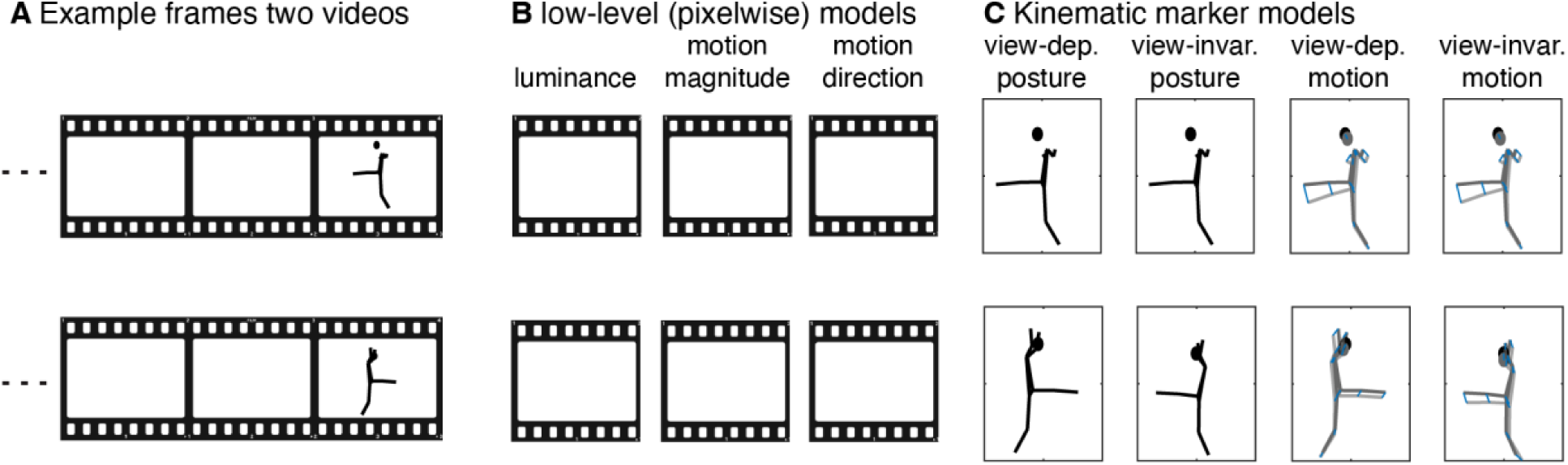
Illustration of stimulus models. Figure adapted from De Vries et al. (2023) Nat Commun, CC BY 4.0 (1). (**A**) Example frames of two videos (rows) to be correlated for creating the model RDMs. (**B**) Low-level visual models. Left: pixelwise luminance (spatially smoothed grayscale). Middle: magnitude of pixelwise motion (operationalized as optical flow vectors), with brighter colors indicating higher magnitude. Right: direction of pixelwise motion, with optical flow vectors indicated in blue and scaled 10 times for illustrative purposes. (**C**) Models based on 3D kinematic marker positions, from left to right: view-dependent body posture, view-invariant body posture (i.e., after aligning the kinematic markers between videos without changing their internal structure, through translation and rotation along the vertical axis), view-dependent body motion as indicated by blue lines (i.e., difference in kinematic marker position between two subsequent frames), and view-invariant body motion. Note that the main dRSA regression-based analysis included also view-dependent and view-invariant acceleration models, as well as gaze-position models based on individual participant eye-tracker data.

**Fig. S5.**
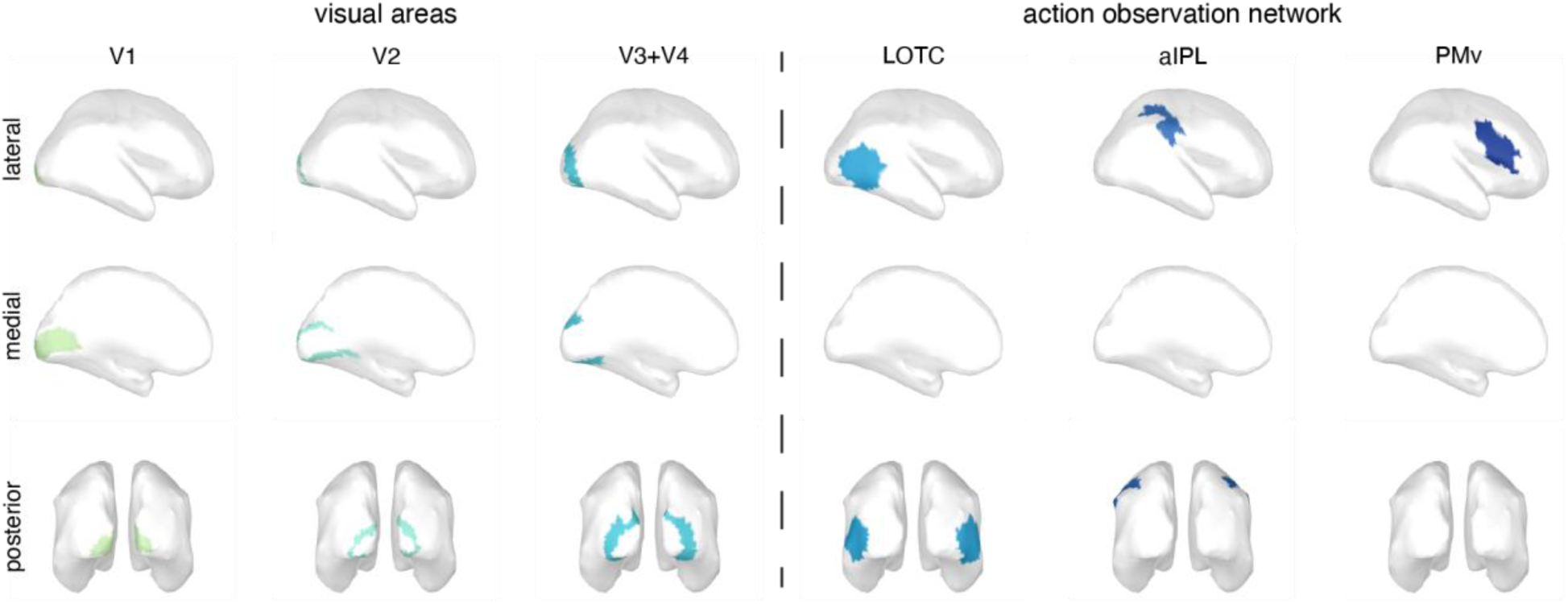
ROI definitions. Figure reproduced from De Vries et al. 2023, Nat Commun, CC BY 4.0 (1). Cortical regions of interest (ROIs) for main analysis based on combinations of parcels as defined according to the Human Connectome Project (HCP) atlas (*2*). ROIs were a priori defined as follows (ROI name = atlas parcels): V1 = V1 [487 vertices]; V2 = V2 [491 vertices]; V3+V4 = V3 and V4 [490 vertices]; LOTC = V4t, FST, MT, MST, LO1, LO2, LO3, PH, PHT, TPOJ2 and TPOJ3 [565 vertices]; aIPL = PF, PFt, AIP and IP2 [403 vertices]; PMv = IFJa, IFJp, 6r, 6v, PEF, IFSp, 44 and 45 [466 vertices]. Note that these vertex amounts are based on the ICBM152 template cortical surface, exact amounts differ slightly between individual subjects. MEG responses at all vertices from both hemispheres were combined into a single vector to compute the neural RDM at a single time point (i.e., pairwise dissimilarity in the MEG response to the 14 action sequences). For visualization, all vertices within a single ROI are given the same color, which matches the color for each ROI in the main results (Fig. 2, 6 and S1). LOTC = lateral occipitotemporal cortex, aIPL = anterior inferior parietal lobe, PMv = ventral premotor cortex.

**Fig. S6.**
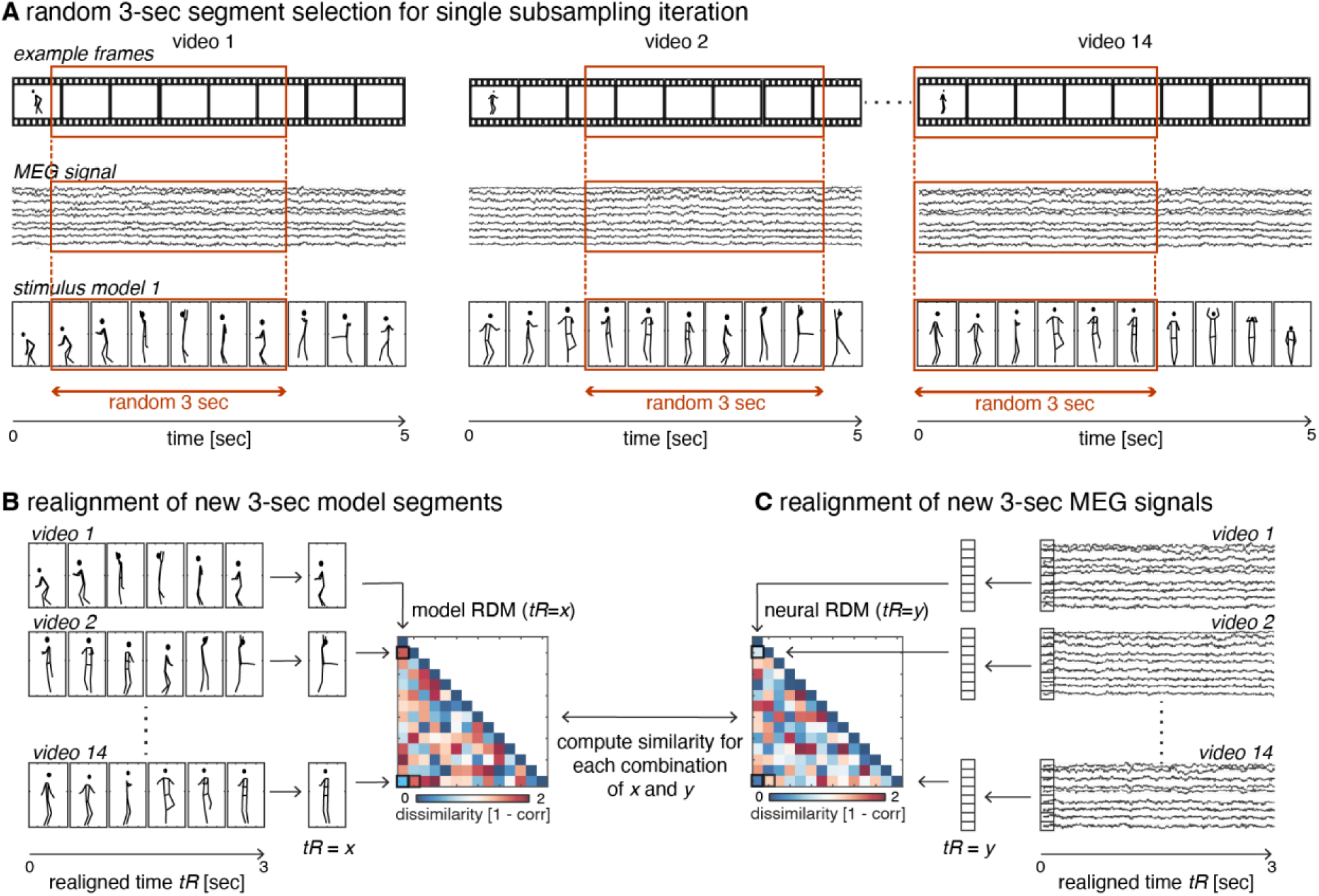
Temporal subsampling and realignment. Figure reproduced from De Vries et al. 2023, Nat Commun, CC BY 4.0 (1). Temporal subsampling and realignment were used to attenuate idiosyncratic temporal heterogeneity in dRSA results caused by arbitrary pairwise alignment specific to these 14 stimuli (see subsection “Temporal subsampling” in the “Methods” section). (**A**) On each of 1000 subsampling iterations, a 3-sec segment is randomly selected independently for each of the 14 stimuli (orange boxes). Crucially, while a different random 3-s window was selected for the 14 stimuli, for a given stimulus the same 3-sec window were selected for both neural and model data, thus keeping temporal alignment between those intact (indicated by vertical orange dotted lines). (**B**) and (**C**) Next, the 14 new 3-s segments are realigned, after which model and neural representational dissimilarity matrices (RDMs) are computed at each realigned time point t_R_. Last, similarity is computed for each combination of realigned model time (x) and neural time (y). Note that the steps in (**A**), (**B**) and (**C**), as well as the last step to compute the dynamic representational similarity analysis (dRSA) curves, are all done within a single subsampling iteration. After 1000 subsampling iterations, dRSA results are averaged over iterations.

**Table S1.**
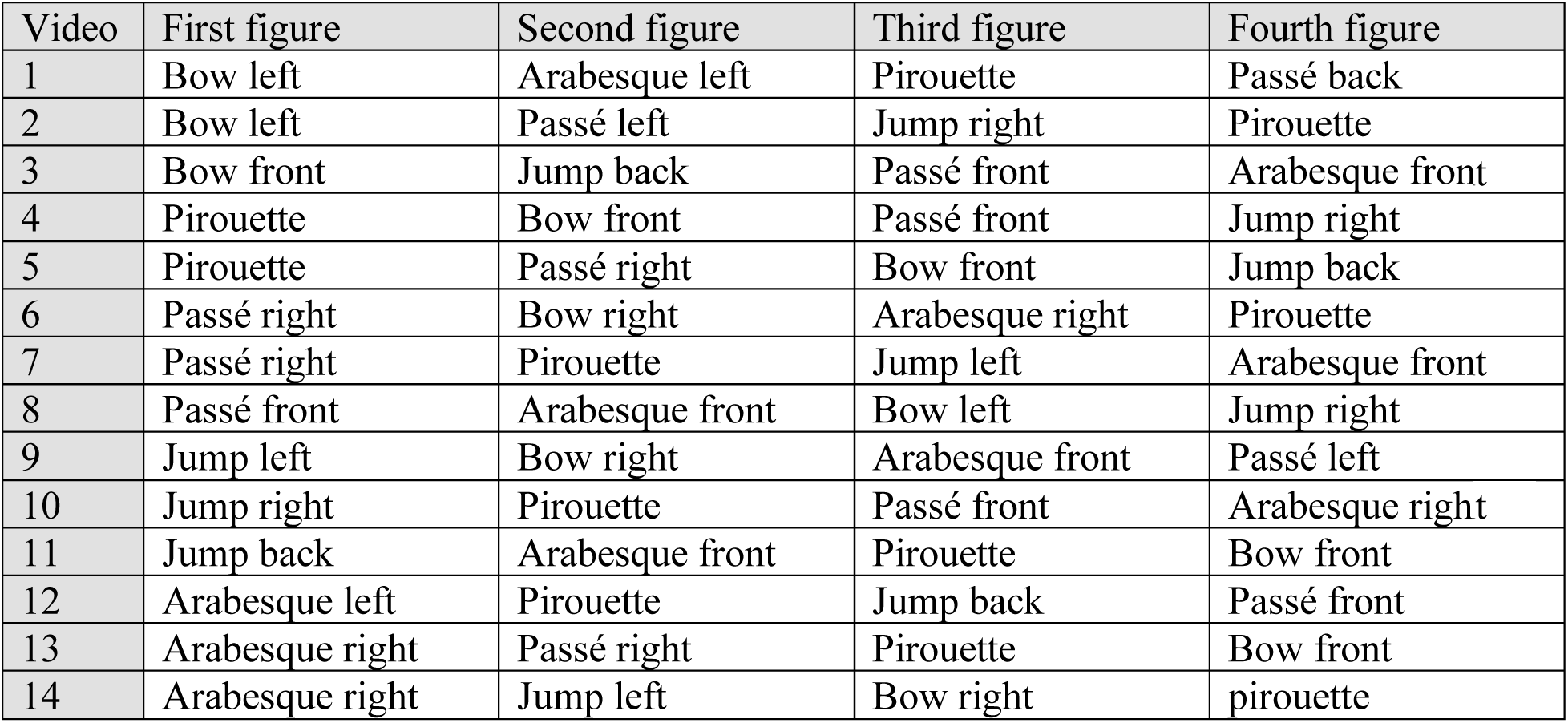
Ballet figures per stimulus. Each video stimulus consisted of a unique sequence of 4 smoothly connected ballet figures. The 4 figures were selected from a total of 5 unique figures. Note that all figures were presented from multiple viewpoints, to allow for both view-dependent and view-invariant body posture and motion models. The pirouette was always performed counterclockwise and could start and end at different angles.

**Table S2.**
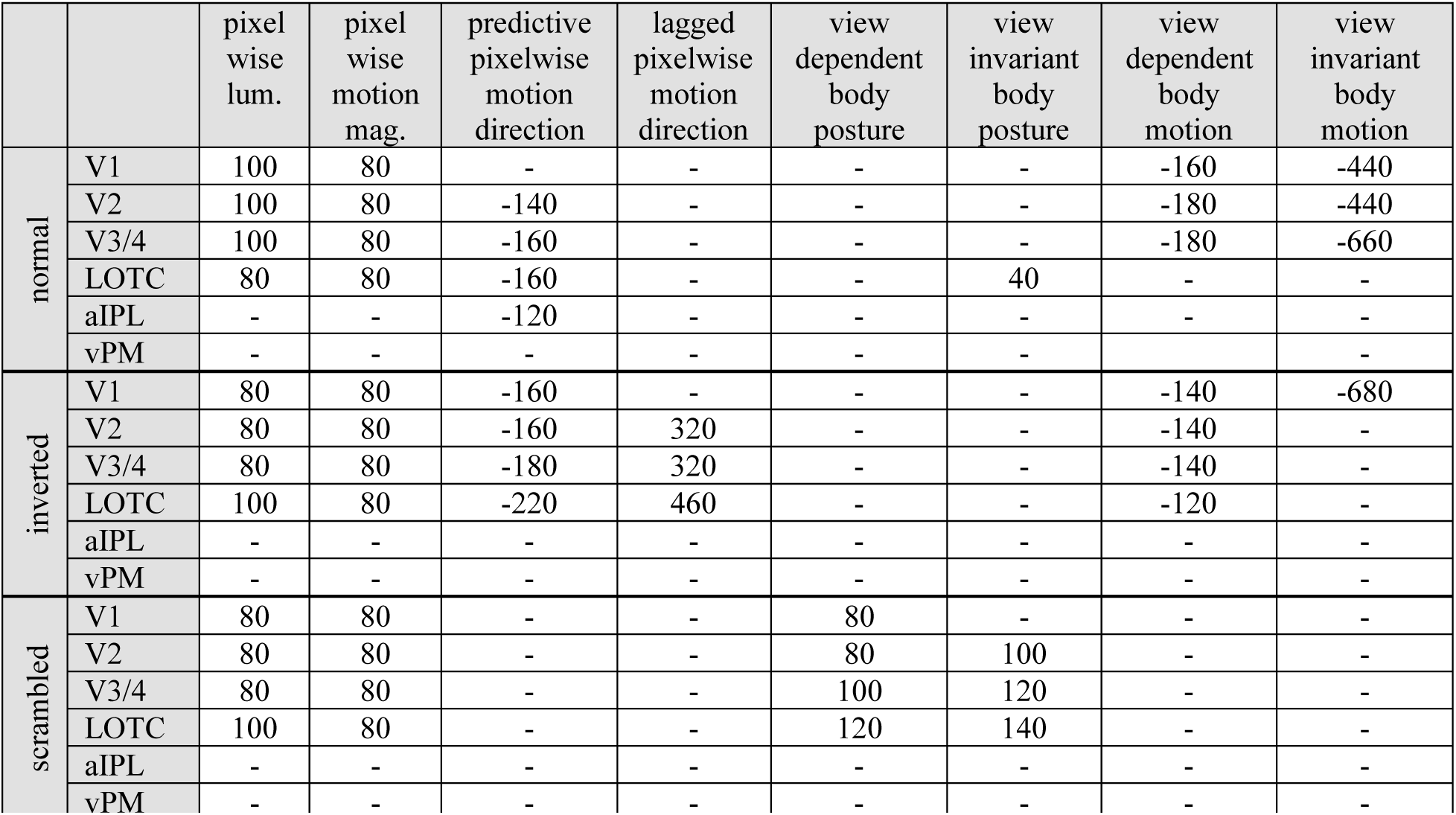
Peak latencies of the participant-average dRSA curves. Peak latencies are indicated in msec relative to zero lag between neural and model RDMs. Results are shown only for condition-ROI-model combinations with a significant main dRSA result (Fig. 2, 6, and S1), since latency estimation is inaccurate otherwise. We observed both a predictive (negative) and lagged (positive) peak for pixelwise motion direction in the inverted condition (Fig. 2b) and therefore computed peak latency separately for each. LOTC = lateral occipitotemporal cortex, aIPL = anterior inferior parietal lobe, PMv = ventral premotor cortex. Lum. = luminance, mag. = magnitude.

## Notes

### Competing Interest Statement

The authors have declared no competing interest.

### Summary of Updates

Informed consent has now been obtained from the ballet dancer for the publication of the videos + example frames used in the article. In this revision this is now clearly stated. Note that this preprint version still contains the placeholder empty frames since BioRxiv does not permit identifiable material, but the final publication will display the original video frames.

